# Aging effects on retrospective and naturalistic encoding of time and their relationship with memory

**DOI:** 10.64898/2026.10.08.757302

**Authors:** Florentine Fricker, Giulia Buzi, Maelys Morantin, Franck Doidy, Patrice Clochon, Raphaël Bordas, Nathan Leroy, Arnaud D’Argembeau, Virginie van Wassenhove, Thomas Hinault

**Affiliations:** Université de Caen Normandie, Inserm, EPHE-PSL, PSL Université Paris, CHU de Caen, GIP Cyceron, U1077, NIMH, 14000 Caen, France; CEA/DRF/Inst. Joliot, NeuroSpin; INSERM, Cognitive Neuroimaging Unit; Université Paris-Saclay, Gif/Yvette, 91191 France; Psychology and Neuroscience of Cognition Research Unit, University of Liège, F.R.S-FNRS, 4000 Liège, Belgium

**Keywords:** Aging, Timing, EEG, Alpha oscillations, Memory

## Abstract

In everyday life, attention is not explicitly directed toward time. Yet, we can reconstruct how long an experience lasted from the memory traces it left behind, a process intertwined with episodic memory and subjective experience. As both memory and temporal processing change with advancing age, investigating aging effects could provide critical insights on these reconstructive processes. In this study, 31 younger (aged 24±3) and 36 older (aged 69±6) healthy adults completed two EEG sessions: 1) a resting-state recording followed by an unexpected retrospective time estimation, and 2) a video-based prospective duration estimation task where participants encoded and judged the duration of 50 naturalistic videos (2-32s) varying in eventfulness. Using a cycle-by-cycle burst analysis of alpha (α: 8-12 Hz), we examined oscillatory correlates of both retrospective temporal judgement and prospective temporal processing. Behaviorally, both groups reconstructed short intervals similarly, but older adults produced longer prospective duration estimates than younger adults for longer video clips (16-32s), particularly under low-eventfulness. Regression analyses also showed that individual differences in older adults’ prospective duration judgments were influenced by both their retrospective time estimation and episodic memory abilities, especially for longer videos. At the neural level, resting-state α-burst dynamics were higher in older adults and predicted, with α-power, longer retrospective estimates in older adults. Additionally, resting-state α-burst activity predicted α-burst during video encoding in both groups. Together, these findings reveal that individual α-power and α-burst dynamics could constitute memory-based time-tracking mechanism that aging reshapes in ways that manifest behaviorally under naturalistic conditions.

## 1. Introduction

In everyday life, time is rarely monitored explicitly. We do not count seconds as they elapse, nor do we consciously track the passage of minutes during a conversation, a meal, or a walk. Yet, when asked how long an experience lasted, or how quickly the past hour seemed to fly by, we can give an estimate drawn from memory traces reconstructed after the fact. This reconstruction is retrospective, incidental, and deeply embedded in the unfolding of experience [1,2]. Despite its centrality in daily life, the cognitive and neural mechanisms underlying such temporal reconstruction remain poorly understood and have been largely overlooked in favor of paradigms requiring explicit prospective attention to time [3], for a review on the lab-dilemna see [4]. Moreover, how these mechanisms operate in richer, naturalistic contexts, and how they are reshaped with advancing age has received little empirical attention [5]. The present study addresses this gap by examining retrospective time estimation and naturalistic duration estimation in younger and older adults, alongside their oscillatory neural correlates.

Every experience holds a temporal dimension and one’s ability to detect and reconstruct various durations is shaped by the richness and structure of ongoing experience [6–9]. Environmental regularities unfold across time; sequence, order, and contextual transitions are necessary to structure experience and anchor it in memory [10,11]. Critically, this temporal structuring is largely incidental, retrieved retrospectively from memory traces formed during ongoing activity [9,12,13]. Recent memory-of-time frameworks further propose that the brain can tag episodic representations with temporal information during encoding, providing a potential substrate for such reconstruction [2,13]. However, temporal judgments have mostly been assessed *prospectively,* that is when attention is directed toward time as it elapses [6,9,14–17]. For example, in bisection tasks, participants have to judge the relative duration of intervals presented successively and indicate whether the second interval was shorter or longer than the first (see [18,19] for a review on timing paradigms). Importantly, temporal judgments can also be made *retrospectively*, that is when time becomes the focus of attention only after it has elapsed and where the request for a duration judgment is unexpected [17,20–22]. Retrospective timing is typically framed within memory-based accounts, according to which duration judgments are reconstructed from stored contextual information [23]. Early storage-size models [24] proposed that perceived duration scales with the amount of information encoded during an interval, later refined into contextual-change models emphasizing transitions in mental context [9], linking retrospective timing to episodic memory processes [25–28]. Memory mechanisms may thus provide the very substrate upon which retrospective time judgments are later reconstructed. However, this temporal reconstruction has been studied almost exclusively using brief intervals with impoverished content that bears little resemblance to the continuous flow of real-life experience, leaving open how it operates when time is embedded in a richer and more dynamic context.

Naturalistic stimuli such as narratives, movies, and virtual reality environments have increasingly been used to study cognition under more ecologically valid conditions [29–31], and time perception is no exception [32–35]. Indeed, probing timing abilities with naturalistic scenes in which stimuli are embedded enhances the temporal context by providing a richer stream of contextual information unfolding over time [36,37]. Importantly, in such contexts, the length and complexity of naturalistic stimuli mean that even prospective duration judgments must draw on memory-based reconstruction, bringing them closer to the reconstructive processes observed in retrospective timing [9,17]. This is particularly the case when durations extend beyond working memory span, such that this temporal reconstruction necessarily relies on the structure of what was experienced rather than on active monitoring [38,39]. Consistent with this view, recent work with naturalistic videos suggests that the retrospective duration of whole events depends primarily on their remembered event structure, with the number of events encoded in memory playing a key role [8], directly linking temporal perception to the way experience is segmented and stored. A key framework for understanding how experience is segmented into memorable units is Event Segmentation Theory (EST; [40,41]), according to which we continuously parse the ongoing stream of experience into discrete events. These events serve as privileged anchors in memory [42–46] and directly shape retrospective duration: the more events encoded (i.e., the level of eventfulness), the longer the interval seems in retrospect [33,47–50]. Eventfulness thus constitutes a key driver of naturalistic duration estimation, with higher eventfulness consistently producing longer estimation relative to low-eventfulness contexts [51,52]. Yet, whether these mechanisms operate similarly across the lifespan remains largely unknown, as most of this work has focused on younger adults.

Indeed, one central factor influencing timing abilities is advancing age. In prospective timing tasks, older adults’ performances often show increased variability and flattened psychometric curves [53–56]. On the other hand, implicit timing abilities appear largely preserved in aging [54,56,57], with retrospective duration judgments showing only little systematic bias [6,37]. Yet, greater inter-individual variability is consistently observed in older adults across these measures, suggesting that preserved mean-level performance may conceal underlying heterogeneity. Since retrospective timing is expected to involve episodic memory processes [25–27] and episodic memory shows the steepest age-related decline [58,59], this variability may be particularly pronounced when temporal reconstruction is most memory dependent. Consistent with this, recent evidence showed that reduced episodic encoding capacity underlies the subjective acceleration of time over the past decade in older adults [5], suggesting that episodic memory resources constitute a common substrate across timescales. However, whether this principle extends to durations ranging from seconds to minutes remains an open question.

With regards to naturalistic timing paradigms, older adults also show altered timing when durations are embedded in semantically rich or realistic visual scenes compared to abstract stimuli, with naturalistic context benefiting younger adults’ precision while impairing older adults’ performance, possibly due to difficulties in suppressing temporally irrelevant contextual information [36,37,60]. As most research on cognitive aging has relied on lab-assessed explicit timing tasks that are far removed from the complexity of daily experience, naturalistic stimuli may provide a better understanding of the factors driving the inter-individual variability frequently observed in cognitive healthy aging [61–64]. Together, these findings suggest that preserved mean-level temporal performance in older adults may conceal underlying neural reorganization that behavioral measures alone cannot capture.

One promising window into age-related neural reorganization comes from resting-state neural oscillations, whose dynamics have been linked to temporal processing across multiple timescales [16,65–67]. Indeed, a slowing of neural oscillations in aging has been consistently observed across frequency bands [68–70]. With its millisecond temporal resolution and sensitivity to gradual oscillatory shifts, electroencephalography (EEG) is particularly well-suited to track the age-related changes in temporal processing and provide insight into the cognitive and neural bases of retrospective temporal judgements across the lifespan. Notably, alpha oscillations (8-12 Hz) could offer a common neural framework, as they have been linked to temporal resolution and episodic structuring more broadly [16]. Recent work suggests that spontaneous alpha burst^1^ dynamics may predict retrospective duration judgments in the absence of explicit timing demands [74,75], consistent with the episodic stamping hypothesis which proposes that alpha bursts passively tag temporal information onto episodic memory traces during ongoing experience. In more immersive contexts, further evidence indicates that distinct spectral patterns may index different temporal experience states [76]. Beyond controlled laboratory paradigms, EEG recorded during naturalistic movie viewing has revealed systematic theta (4-7 Hz), alpha, and beta (13-30 Hz) activity in fronto-parieto-occipital regions linked to event segmentation operations [77], suggesting that oscillatory dynamics may track the mechanisms that structure naturalistic temporal experience. However, whether these signatures, and especially alpha oscillations, generalize to naturalistic duration estimation paradigms, and how they are modulated by age, remains an open question. Critically, these oscillatory signatures are known to be sensitive to the richness and structure of the encoded context [75], making them particularly relevant for probing temporal reconstruction as addressed in the present study.

Taken together, these observations converge on a potential central role of alpha oscillations and episodic memory changes with advancing age to retrospective time judgments and naturalistic duration estimates. Here, we investigate these changes at both behavioral and oscillatory levels. First, we characterize retrospective time estimation alongside its oscillatory correlates, in particular with regard to alpha band activity, in younger and older adults. Second, we characterize naturalistic duration estimation across durations and eventfulness levels and similarly investigate its oscillatory correlates and its relationship with retrospective timing and age. Third, we investigate the extent to which episodic memory resources constitute a common substrate linking these different forms of temporal judgment, and how this relationship is modulated by age. By bridging retrospective timing and naturalistic duration estimation within a lifespan perspective, this study aims to clarify how the brain reconstructs temporal experience, and how these mechanisms are reshaped with age.

## 2. Method

### 2.1. Participants

Seventy-one French-speaking volunteers participated in the experiment. All participants reported normal or corrected-to-normal vision and hearing, and no history of major medical conditions, neurological or psychiatric disorders, or head trauma. Older adults were screened using the Montreal Cognitive Assessment (MoCA; [78]) and were included only if they scored ≥ 26 (mean score = 28.39 ± 1.18, ranging from 26 to 30).

The study was approved by a French ethics committee (Comité de Protection des Personnes, CPP; ID-RCB: 2024-A00167-40), and all participants provided oral and written informed consent in accordance with the Declaration of Helsinki (2024). Participants were recruited through academic mailing lists and social media and received 20 € for their participation.

Four participants were excluded from the final analyses: two due to excessive EEG noise, one because participant reported anticipating the retrospective duration judgment during the first session, and one due to a technical issue during acquisition.

The final sample consisted of 67 participants: 31 younger adults (17 females; mean age = 23.81 ± 2.97 years, ranging from 20 to 31 years old) and 36 older adults (26 females; mean age = 69.06 ± 5.60 years, ranging from 61 to 79 years old). Additionally, two participants (one younger adult and one older adult) did not complete the second session and were consequently excluded from analyses involving second session tasks but were retained for first-session analyses.

### 2.2. Procedure

Participants completed two experimental sessions separated by approximately one month (mean interval = 30.28 days, min = 18, max = 61), as part of a larger research project. Each session lasted approximately three hours. Retrospective timing tasks were run on Opensesame (version 4.1.7; [79]) and the video-based duration estimation task was run on E-Prime 3.0 (Psychology Software Tools, Pittsburgh, PA).

#### Session 1 (T1)

During the first session, older adults were screened using the MoCA to confirm eligibility. After EEG net installation, participants were seated approximately 90 cm from a computer screen inside a Faraday cage and underwent an open-eyed resting-state EEG recording corresponding to the retrospective timing task (where rTE1, retrospective time estimate ratio at T1, is collected). Then, they completed the retrospective post-task questionnaire and the first part of neuropsychological assessment battery.

#### Session 2 (T2)

During the second session, the researcher explained the video-based duration estimation task to the participants and provided them with written instructions. Participants then started a training block composed of four trials. The training was repeated if requested by the participant. Data from the training session were not included in the analyses.

After EEG net installation, participants first underwent a second resting-state EEG recording including the retrospective timing task (where rTE2, retrospective time estimate ratio at T2, is collected), followed by the video-based duration estimation task and its self-report questionnaire. Finally, they completed the remaining part of neuropsychological assessment battery. The overall experimental timeline is illustrated in Fig. 1a.

**Fig. 1.**
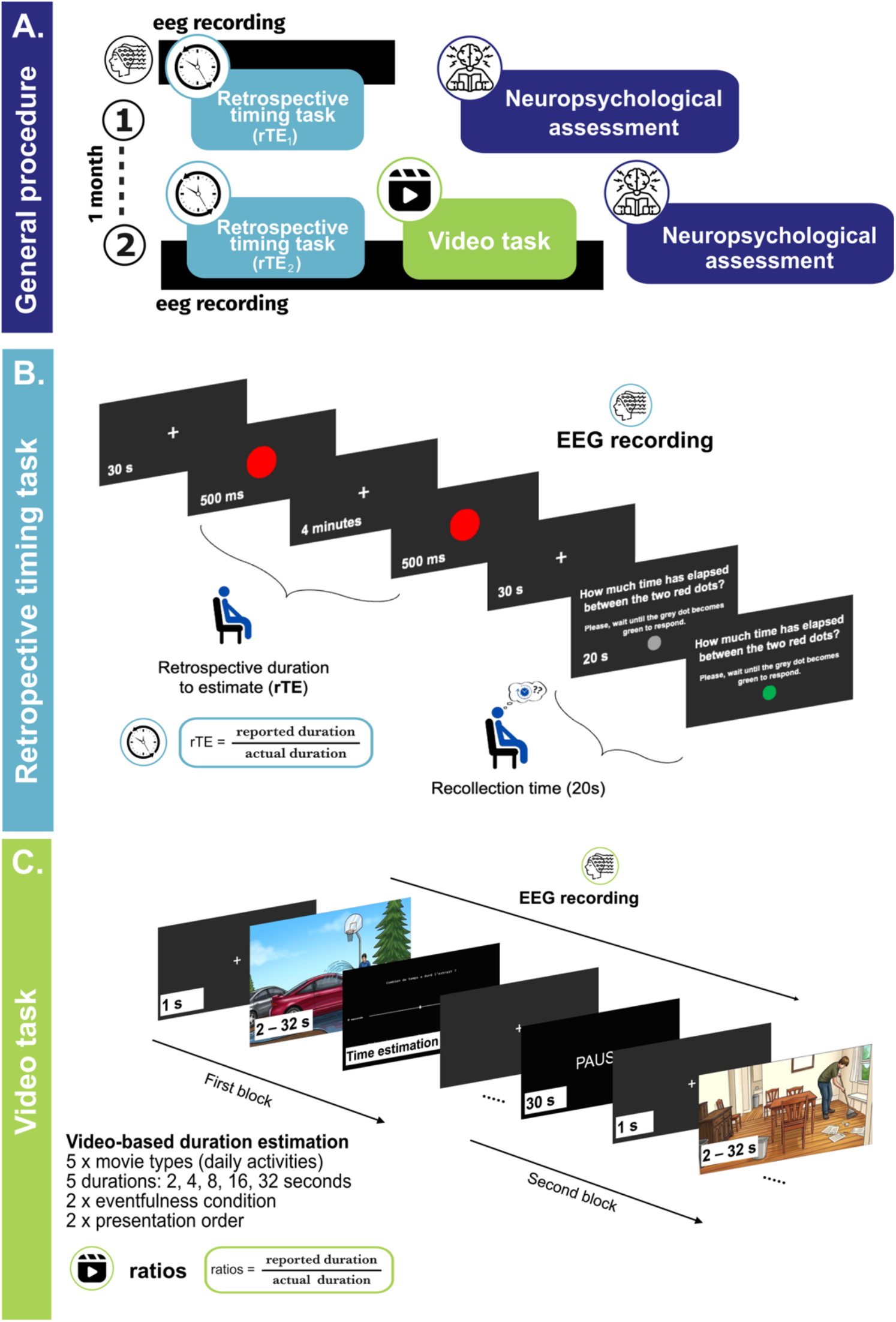
General experimental procedure sessions and overview of the retrospective timing and the video-based prospective duration estimation tasks. a) *General procedure.* Participants completed two sessions separated by approximately one month. Session 1 comprised resting-state EEG recording including the retrospective timing task (where rTE1 is collected), its post task questionnaire and completion of the first part of the neuropsychological battery. Session 2 included a second resting-state EEG recording with the retrospective timing task (where rTE2 is collected), the video-based duration estimation task and its self-report questionnaire, and completion of the remaining neuropsychological assessments. b) *Retrospective timing task*. Participants remained in quiet wakefulness during a 4-min EEG recording. No information about a subsequent time judgment was provided. At the end of the recording, they were asked to estimate as precisely as possible the duration that had elapsed between the two red dots marking the beginning and end of the interval. For each participant, retrospective time estimates were expressed as the ratio between the verbal report and the objective clock duration, separately for the first (rTE1) and second (rTE2) sessions. c) *Video-based prospective duration estimation task.* Participants performed a video-based prospective duration estimation task while undergoing EEG recording. Each trial began with a fixation cross (1 s), a video clip (2-32 s), and a response screen where participants had to indicate their duration estimate thank to a non-graduated slider (0-45 s). The design included 50 trials (5 movie types x 5 durations, x 2 eventfulness levels) presented in two blocks. *Illustrations of the video content were generated with artificial intelligence and replace the original video snapshots in this preprint version*.

### 2.3. Experimental tasks

#### 2.3.1. Retrospective timing task at session 1 and 2

Participants underwent an open-eyed resting-state EEG recording paradigm adapted from the quiet wakefulness retrospective time paradigm described in previous studies [74,75]. Before the recording, the experimenter instructed participants that their brain activity would be recorded at rest. Critically, they were not informed that they would later be asked to estimate time, nor that the study concerned time perception. They were asked to remain still, keep their eyes open, fixate a central cross on a screen in front of them, and to “clear their mind” throughout the recording.

Following these instructions, the experimenter initiated the EEG recording. From the participant’s perspective, the sequence unfolded as follows (see Fig. 1b): a fixation cross appeared on the screen. Thirty seconds after the beginning of the recording, a red dot was presented centrally for 500 ms. The fixation cross then reappeared and remained on the screen. Four minutes after the first red dot disappearance, a second red dot appeared for 500 ms, thereby marking the temporal boundaries of the episode. 30 seconds after the second red dot disappearance, participants were asked to estimate as precisely as possible the duration that had elapsed between the two red dots. A grey dot was displayed for 20 seconds before turning green, signaling that participants should wait for the experimenter to enter the room before providing their verbal response. They were instructed to report their estimate as precisely as possible, using minutes and seconds and avoiding approximations.

To ensure that participants were unaware of the timing task, the experimenter asked right after whether they had anticipated this question. Because this paradigm was administered at both the first and at the second session one month later, this control was essential to ensure the task constituted a proper retrospective timing paradigm, in which duration estimates rely on post hoc reconstruction of the encoded episode rather than on prospective timing processes. One participant (rejected from the final dataset) expected the question in the first session; 23 younger adults (i.e., 76.7 %) vs. 9 older adults (27.3 %) expected the question in the second session (χ^2^(1) = 15.34, *p* < .001).

For analysis, participants’ retrospective time estimates at the first (rTE1) and second (rTE2) sessions were expressed as the ratio of individual’s verbal report to the objective clock time (four minutes). Ratios greater than 1 indicate overestimation, while ratios below 1 indicate underestimation.

##### Post-task questionnaire

At the first session, participants also completed a questionnaire adapted from Wang and collaborators [83] designed to assess their subjective experience during the resting-state recording. Questions comprised items rated on a 5-point Likert scale and related to mental orientation (e.g., temporal, spatial and social orientation), thoughts content (e.g., internal dialog, vividness, volition, valence, density, self-orientation and richness), feelings (e.g., patience, entertainment, boredom, fatigue, happiness, relaxation, calm, preoccupation), and passage-of-time judgment expressed on a 5-point Likert scale, going from “very slow” to “very fast” (see Supplementary Fig. S1 for more details and results).

#### 2.3.2. Video-based duration estimation task

During the second session and after the retrospective timing task, participants performed a video-based prospective duration estimation task while undergoing EEG recording. Participants were explicitly informed before this task that they would need to estimate the duration of each clip. The task consisted of 50 trials organized into two blocks as follows (see Fig. 1c): each trial began with a fixation cross displayed on the screen for one second, followed by a pseudo-randomized presentation of a video clip lasting either 2, 4, 8, 16, or 32 seconds. For each trial, participants estimated the duration (in seconds) of the clip using a slider controlled by the computer mouse answering with their right hand. The slider ranged from 0 to 45 seconds in one-decimal increments, with the minimum (0 second) and maximum (45 seconds) values displayed at each end of the scale. The cursor was positioned at the midpoint of the scale to discourage counting strategies, and the scale was not graduated to avoid anchoring to standard values. Participants responded without time constraints and were allowed a 30-second break between blocks.

##### Behavioral data

For each trial, the duration judgment was expressed as the ratio of the participant’s duration estimate to the objective clip duration. Again, ratios greater than 1 indicate overestimation, while ratios below 1 indicate underestimation. Mean ratios were then computed for each duration-eventfulness combination.

##### Experimental stimuli

Video clips were extracted from five longer movies commonly used in event segmentation studies (i.e., [80–82]), depicting everyday activities performed by a single actor. The original movies were filmed from a fixed viewpoint at approximately standing head height. Cinematic effects such as cuts, pans, and zooms were not used to avoid introducing external segmentation cues. Similarly, all clips were presented without sound to avoid additional temporal cues through auditory information. Given the influence of emotional valence and arousal on time perception [84–86], only emotionally neutral scenes were used. Five movie types were selected for the main experiment (*Binder*, *Breakfast*, *Car*, *Sweeping*, *Tire*) depicting common activities (e.g., photocopying a book, preparing breakfast, washing a car, sweeping, changing a tire). For the training session, three additional movies depicting similar everyday activities were used. Participants were seated approximately 90 cm from the computer screen, and each movie was displayed full screen on a 47.6 cm x 26.8 cm monitor (resolution 1920 × 1080).

##### Video clips extraction and creation

Event segmentation data were obtained from previous studies [80–82], in which participants were instructed to press a button whenever they perceived an event boundary, defined as the moment when one meaningful unit of activity ended and another began. Following a procedure adapted from Bangert and collaborators [47], video excerpts were selected in three steps. First, for each movie, each second was assigned an event score corresponding to the number of participants who pressed the button during that second. Second, for each movie and each possible temporal window matching our target durations (2, 4, 8, 16, and 32 seconds), the mean event score per second was computed. Third, excerpts were selected to match either the maximum (high eventfulness) or minimum (low eventfulness) mean event score observed across all possible 32-second windows, yielding two eventfulness conditions (high: M = 6 keypresses/s; low: M = 1 keypress/s). Excerpts were then extracted using iMovie (v10.4.4) at five durations (2, 4, 8, 16, and 32 seconds), ensuring no temporal overlap (separated by minimum 2 seconds, 20 seconds on average). This resulted in 50 unique video clips (5 movie types x 2 eventfulness conditions x 5 durations). The complete list of video clips and their features is provided in Supplementary Fig. S2.

##### Randomization

Regarding presentation order, two fixed-order lists (AB and BA), each comprising two blocks, were used and counterbalanced across participants. Block durations, corresponding to the cumulative video playback time excluding participants’ response times, were equalized (Block A: 4 min 56 s; Block B: 4 min 58 s; total: 9 min 54 s). Within blocks, trials were pseudo-randomized such that no movie type appeared twice consecutively and each type was approximately equally represented across blocks.

##### Post-task questionnaire

Immediately after completion of the video task, participants filled out a brief self-report questionnaire assessing their subjective experience during the task. Questions concerned perceived confidence, task difficulty, emotional state (happiness, anger, calmness, sadness), rated on a 5-point Likert scale. Two open-ended questions assessed for task remarks and estimation strategies used (see Supplementary Fig. S3 for more details and results).

#### 2.3.3. Neuropsychological Assessment and questionnaires

Participants additionally underwent a comprehensive neuropsychological and questionnaire-based assessment to characterize cognitive functioning across domains (see Table 1). Socioeconomic status was assessed with the French adaptation of the Hollingshead’s four-factor Index of Socio-Economic status [87].

**Table 1.**
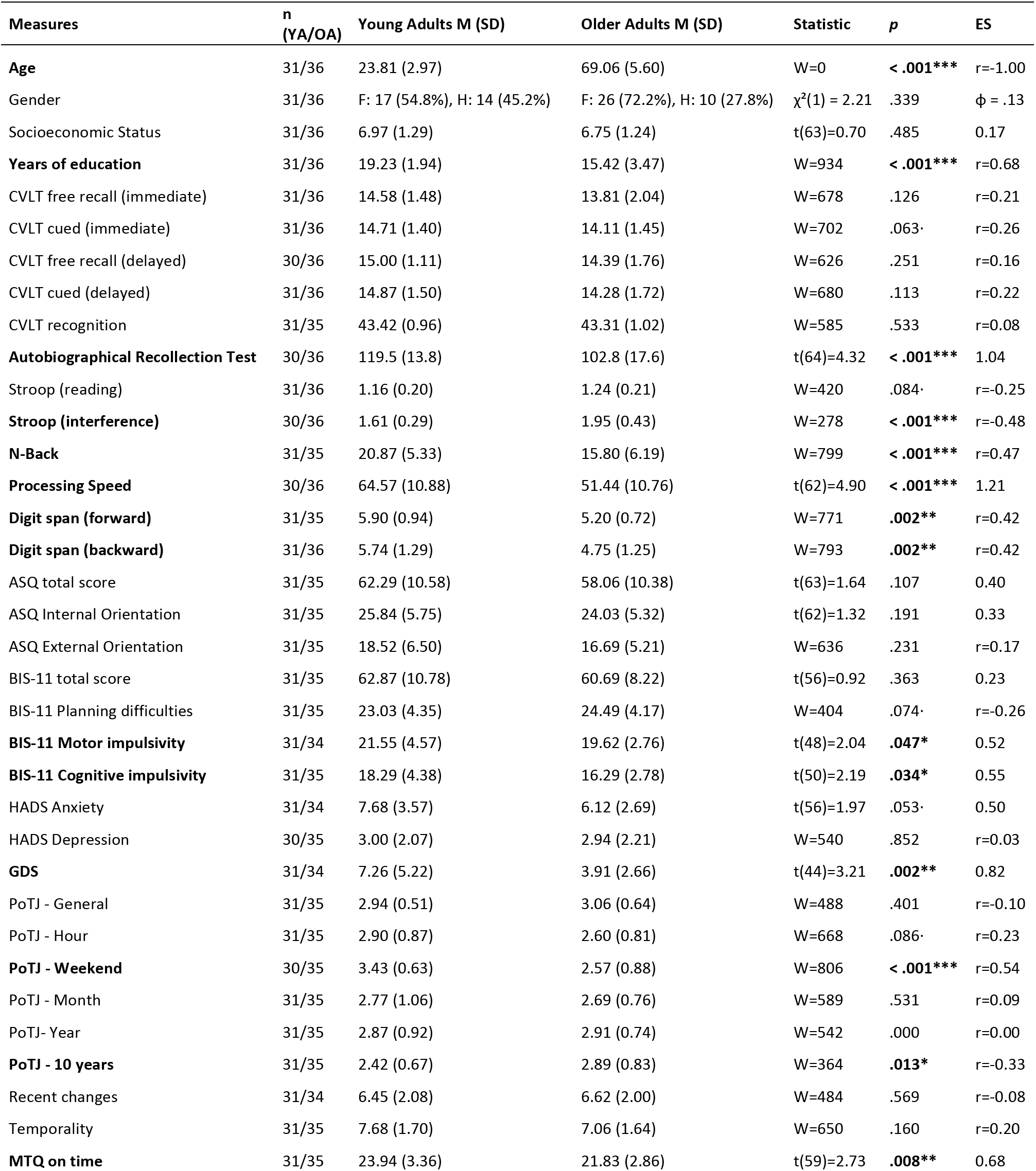
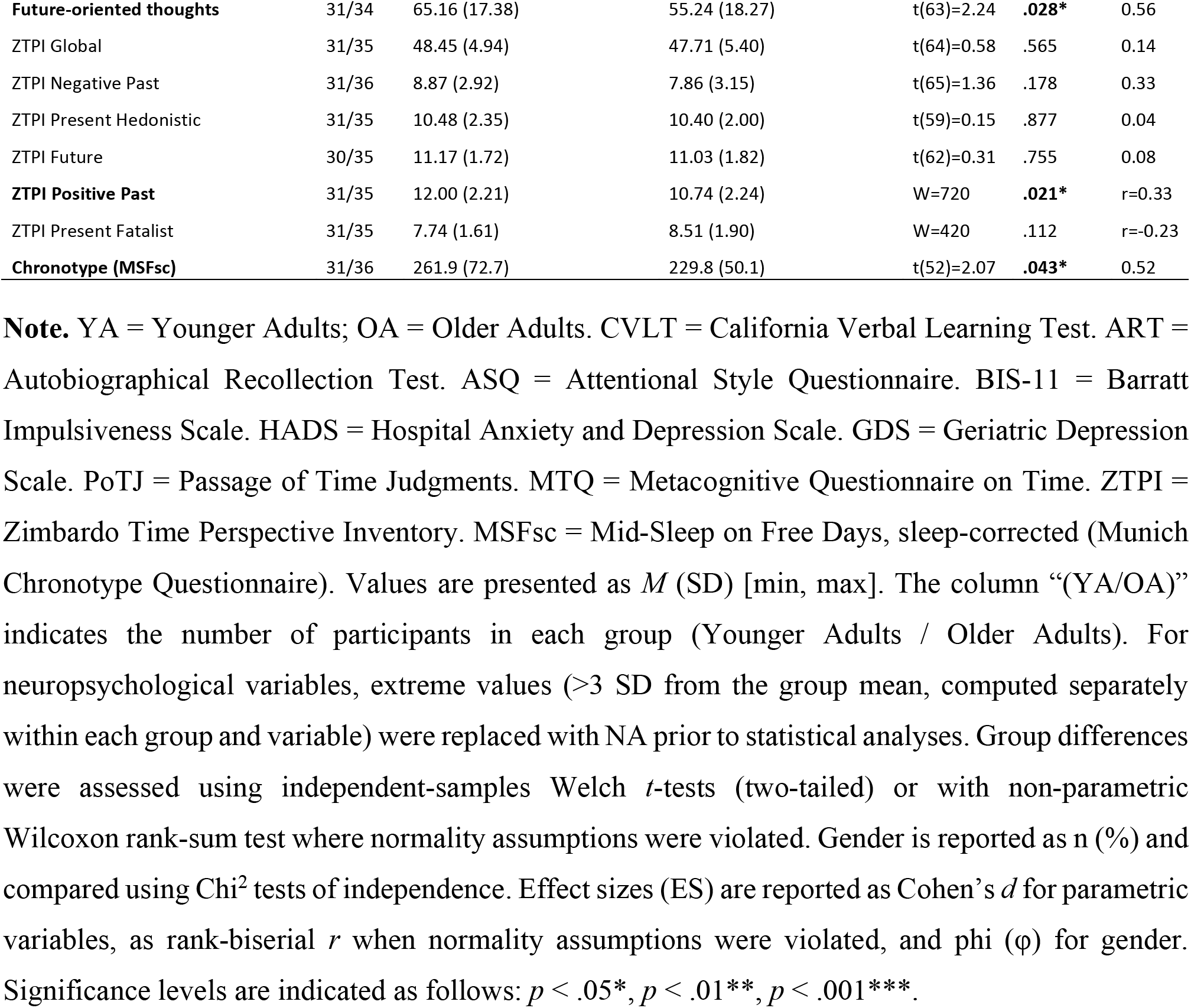
Descriptive statistics and group comparisons for all neuropsychological and questionnaire measures.

Episodic memory was assessed using the California Verbal Learning Test (CVLT; [88]). Autobiographical memory was measured using the Autobiographical Recollection Test (ART; [89]), which captures individual differences in the subjective richness and vividness of recollective experience. Executive functioning was evaluated through inhibitory control using the Victoria Stroop Test [90,91] and working memory using both an N-Back task [92] and digit span measures from the Wechsler Adult Intelligence Scale (WAIS-III; [93]). Processing speed was assessed with the Digit Symbol Substitution Test (WAIS-III; [93]). Attentional orientation toward internal versus external focus was measured using the Attentional Style Questionnaire (ASQ; [94]). Trait impulsivity was assessed with the Barratt Impulsiveness Scale (BIS-11; [95,96]). Depression and anxiety were measured using the Hospital Anxiety and Depression Scale (HADS; [97]) and the Geriatric Depression Scale (GDS; [98]).

Given the focus on temporal cognition, participants also completed several questionnaires probing subjective temporal experience. Passage-of-time judgments on longer scales (general, next hour, weekend, year, ten years) and perceived recent changes were assessed using Wittmann & Lehnhoff’s [99] time experience questionnaire. Metacognitive beliefs about time were further evaluated using the Metacognitive Questionnaire on Time [100]. Temporal perspective was measured using the short version of the Zimbardo Time Perspective Inventory (ZTPI; [101], french adaptation: [102]) capturing negative past, positive past, present hedonistic, present fatalistic, and future orientations. Chronotype was assessed with the ultra-short Munich Chronotype Questionnaire µMCTQ; [103,104]).

### 2.4. EEG recording and preprocessing

#### EEG acquisition

EEG signal was recorded continuously in a faraday cage [105] (GES 400 amplifier, EGI Hydrocel Geodesic Sensor Net, 128 Ag/AgCl dense array sensors), referenced (vertex reference Cz, ground reference CPz), and sampled at 1 kHz frequency with a 24-bit A/D. Impedances were kept under 100 kΩ, with a majority of electrodes below 50 kΩ [106]. Horizontal and vertical electrooculograms (EOG) were recorded during the session to help in artifacts cleaning.

#### EEG Preprocessing

EEG raw data were processed offline using Brainstorm (version 3.260407; Tadel et al., 2011) running under Matlab environment (Version: 24.1.0.2689473 (R2024a) Update 6; The MathWorks Inc., 2024). Data were band-pass filtered between 0.1 and 60 Hz and notch filtered at 50 Hz. Ocular and cardiac artifacts were detected and corrected using signal-space projection (SSP). The number of removed SSP components was comparable across recordings, with a mean of 3.80 (range: 2-8) for the retrospective timing task at T1, 4.16 (range: 1-9) for the retrospective timing task at T2, and 3.39 (range: 1-7) for the video-based duration estimation task. Data were visually inspected, and noisy segments were rejected. Subjects were retained only if at least 75% of the recording remained artifact-free after rejection. EEG data were then re-referenced using the algebraic average of the left and right mastoid electrodes [108]. Bad channels were identified and interpolated using spherical spline interpolation. On average, 5.68 ± 3.93 (retrospective timing task at T1), 5.21 ± 3.39 (retrospective timing task at T2), and 1.83 ± 2.71 (video task) channels were interpolated per participant (4.43%, 4.07%, and 1.43% of the 128 scalp electrodes, respectively).

#### Epoching

For the retrospective timing tasks, EEG data were then extracted time-locked to the two red dots (−100 ms before the first red dot onset to 100 ms after the second red dot onset).

For the video-based duration estimation task, EEG data were segmented time-locked to stimulus onset from −400 ms before each video onset to the end of each clip (2, 4, 8, 16, and 32 seconds). Baseline correction was applied using the pre-stimulus interval (−400 to 0 ms). For each of the 10 duration x eventfulness conditions, the five epochs corresponding to different movie types were averaged, yielding one mean epoch per condition per participant. No epochs were rejected at the trial level; artifact rejection was performed at the continuous signal level, with on average 96.3 ± 3.3% of the recording retained (range: 80.3-98.6%).

#### Spontaneous oscillatory spectrum and localizers

For the retrospective timing tasks, the following analyses were conducted using MNE Python [109] and followed the procedure described in [75], adapted to the present dataset. A brief summary is provided below.

First, to detect alpha activity (8-12 Hz) at each channel, the oscillatory and aperiodic (1/f slope) components of the power spectrum were dissociated using a custom Python implementation of the IRASA algorithm [110]. Continuous EEG data were segmented into fixed-length 5-s epochs, and power spectral density was computed using Welch’s method (1-45 Hz). The slope of the aperiodic component was estimated in log-log space following [111] to control for broadband spectral variations and subtracted to obtain the oscillatory spectrum. Alpha power was then defined as the average oscillatory power between 8 and 12 Hz.

Cluster-based permutation *t*-test (CBPT, [112]) was then performed to identify channels exhibiting significantly elevated alpha power relative to the global average. Preliminary CBPTs conducted separately across age groups and sessions revealed comparable topographies; recordings from Sessions 1 and 2 were therefore pooled across age groups (N = 134 observations) to enhance statistical power and robustness. Clusters were defined using spatial adjacency (Delaunay triangulation implemented in MNE-Python) and 4,000 permutations (Monte Carlo approximation), with a cluster-forming threshold corresponding to the 99.9th percentile of the *t* distribution (*q*(134) = 2.35). The *p* value was estimated based on the proportion of permutations exceeding the observed maximum cluster-level test statistic. Only clusters with corrected *p* < 0.05 are reported. The resulting cluster was consistently located over occipito-parietal channels (Fig. 2a), in line with previous findings. Importantly, this procedure served only to define regions of interest and was not used for statistical inference.

**Fig. 2.**
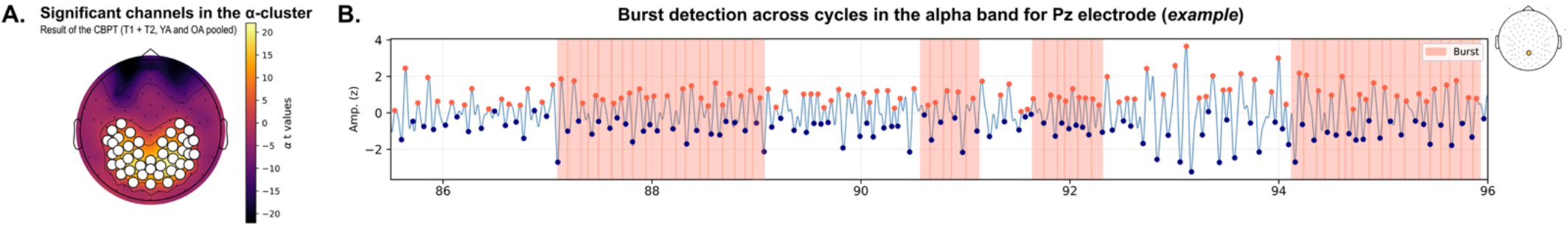
Identification of alpha cluster in the retrospective tasks and burst detection example. a) Occipito-parietal alpha (α: 8-12 Hz) cluster identified using a cluster-based permutation test (4,000 permutations) on oscillatory power following IRASA-based separation of periodic and aperiodic components. To increase statistical power for cluster definition, EEG recordings from both sessions (retrospective timing tasks at T1 and T2) and both age groups (Younger and Older adults) were pooled (total N = 134 observations). b) Illustration of cycle-by-cycle burst detection (bycycle toolbox, [71]) on an example 10-second segment of alpha-filtered EEG signal at electrode Pz, starting 86 seconds after event onset (the first red dot). Peaks (orange dots) and troughs (navy dots) mark individual oscillatory cycles; cycles meeting predefined amplitude, period, and monotonicity thresholds are classified as burst cycles and highlighted in red. Relative burst time corresponds to the percentage of cycles classified as burst cycles across the full recording.

The individual alpha peak frequency (iAPF), computed on the epoched resting-state recordings of the retrospective timing task at both T1 and T2, was defined as the frequency corresponding to the local maximum of the oscillatory spectrum within the 8-12 Hz range and averaged across significant channels in the alpha cluster on a per individual basis.

#### Cycle-by-cycle analysis and burst detection

Bursting features were quantified using the bycycle toolbox [71]. Burst detection parameters were selected to match previously validated settings in similar resting-state paradigms ([74,75]: amplitude fraction threshold = 0.2; amplitude consistency threshold = 0.4; period consistency threshold = 0.4, monotonicity threshold = 0.8, minimum number of contiguous cycles = 3).

For alpha bursts, and prior to the thresholding algorithm, signals were bandpass filtered between 4 and 16 Hz to avoid edge effects related to narrow transition bands. Burst cycles with a frequency outside the 8-12 Hz range were discarded unless they were flanked by burst cycles and the mean burst period fell within an extended alpha range (7.5-12.5 Hz), preventing artificial burst fragmentation due to cycle-to-cycle variability.

Burst detection was applied to two distinct tasks. For the retrospective timing tasks, burst detection was performed on continuous resting-state EEG recordings from retrospective tasks across all 128 channels, using electrode cluster identified from CBPT result as channels of interest. For the video-based duration estimation task, alpha burst detection was applied to averaged EEG epochs time-locked to video onset, separately for each combination of eventfulness (high, low) and duration (2, 4, 8, 16, 32 seconds), yielding 10 conditions per participant.

The primary outcome metric was relative burst time (expressed in percentage), defined as the proportion of cycles belonging to a burst relative to the total number of cycles in the signal (Fig. 2b). Higher values indicate more sustained rhythmic activity. Additional burst features (e.g., number of cycles, mean voltage, mean duration, rise-decay symmetry, etc.) were extracted for alpha cluster as well as across all channels. More details on these metrics are reported in Supplementary Table S1.

### Statistical analyses

Statistical analyses were conducted in R (version 4.5.2; R Core Team, 2025-09-02, ucrt build, https://cran.r-project.org). Missing values were excluded at the variable level, while participants were retained unless all observations for a given analysis were missing. Observations exceeding ±3 standard deviations from the group mean were treated as outliers and excluded prior to inferential testing. Assumptions of normality and homogeneity of variance were assessed using Shapiro-Wilk and Levene’s tests, respectively before statistical analyses were applied.

Unless indicated otherwise, all results presented below are means ± standard deviation (SD). Due to significant differences between group, years of education were included as a covariate in all between-group analyses and partial correlations, which were conducted separately for each group using Pearson or Spearman methods based on normality assumptions. Its influence is reported only when significant. To control for multiple comparisons, *p*-values were adjusted using the Benjamini-Hochberg false discovery rate (FDR) procedure within each family of tests. Results were considered statistically significant at the *p* ≤ 0.05 level.

#### Retrospective encoding of time

Age group-related differences in retrospective time estimates (rTE1, rTE2) were assessed using ANCOVAs with Group (Young, Older) as the between-subject factor and years of education as a covariate. For rTE2, expectancy (whether participants anticipated the timing question) was additionally included as a covariate given its potential influence on retrospective duration estimates. A mixed-design ANCOVA with Session (T1, T2) as within-subject factor was used to examine expectation effects on rTEs across sessions. To test for under-or overestimation, rTEs were compared against veridical relative duration (i.e., 1) using one-sample *t*-tests, or Wilcoxon signed-rank tests when normality was violated.

Age group-related differences in resting-state EEG metrics (iAPF, alpha power, alpha burst activity) were assessed using independent-samples Welch *t-*tests when normality assumptions were met, and Wilcoxon rank-sum tests otherwise. Partial correlations between temporal measures (rTE1, rTE2) and resting-state EEG metrics additionally controlled for expectancy for rTE2. Effect sizes are reported as Cohen’s *d* for parametric tests and rank-biserial *r* for non-parametric tests.

#### Naturalistic prospective encoding of time

Naturalistic prospective time estimation was assessed using a mixed-design ANOVA with Group (Young, Older) as the between-subject factor and Duration (2, 4, 8, 16, 32 seconds; treated as a categorical variable), Eventfulness (low, high) as within-subject factors, and years of education as a between-subject covariate. The same design was applied to alpha burst dynamics during the video-based duration estimation task.

Associations between alpha burst dynamics during the video-based duration estimation task and naturalistic duration estimation ratios were tested diagonally, pairing alpha burst activity and estimation ratio within the same duration and eventfulness condition.

#### Associations between retrospective and naturalistic prospective encoding of time

Partial correlations between temporal measures (rTE1, rTE2) and naturalistic duration ratios were assessed with FDR correction applied within each temporal measure across long-duration clips only (16-s and 32-s, both eventfulness levels), yielding families of 4 tests per temporal measure per group. This restriction was motivated both theoretically, as durations exceeding working memory span rely more heavily on retrospective memory reconstruction [38,39], and empirically, as group differences in naturalistic estimation were specific to long clips in the present data. Two hierarchical regressions were then conducted with the 32-s high- and low-eventfulness duration ratio as the dependent variable.

Correlations between resting-state burst dynamics at T1 and T2 and burst dynamics during the video-based duration estimation task were assessed with FDR correction applied within each band across durations.

## 3. Results

### 3.1. Retrospective encoding of time

#### Similar retrospective time estimates between groups and a general tendency to overestimate the duration of the second session

At T1, both groups underestimated the retrospective duration (M_young_ = 0.81 ± 0.28, *t*(29) = - 3.66, *p* = .001, *d* = −0.67; M_old_ = 0.74 ± 0.34, *t*(34) = −4.63, *p* < .001, *d* = −0.78), whereas estimates were closer to the actual duration at T2 (*p*s > .829; M_young_ = 1.07 ± 0.47; M_old_ = 1.07 ± 0.70). No significant group difference was observed for rTE1 (*β* = −0.42, *t*(63) = −0.43, *p* = .666) or rTE2 (*β* = 0.30, *t*(62) = 1.51, *p* = .137), even after controlling for years of education and, for rTE2, expectancy (Fig. 3a). Years of education significantly predicted rTE2 independently of group (*β* = 0.69, *t*(62) = 2.65, *p* = .010), indicating that participant with more education overestimated the duration at T2. No significant difference in inter-individual variability was observed between groups for either rTE1 (*F*(1, 63) = 0.84, *p* = .363) or rTE2 (*F*(1, 62) = 3.13, *p* = .082).

**Fig. 3.**
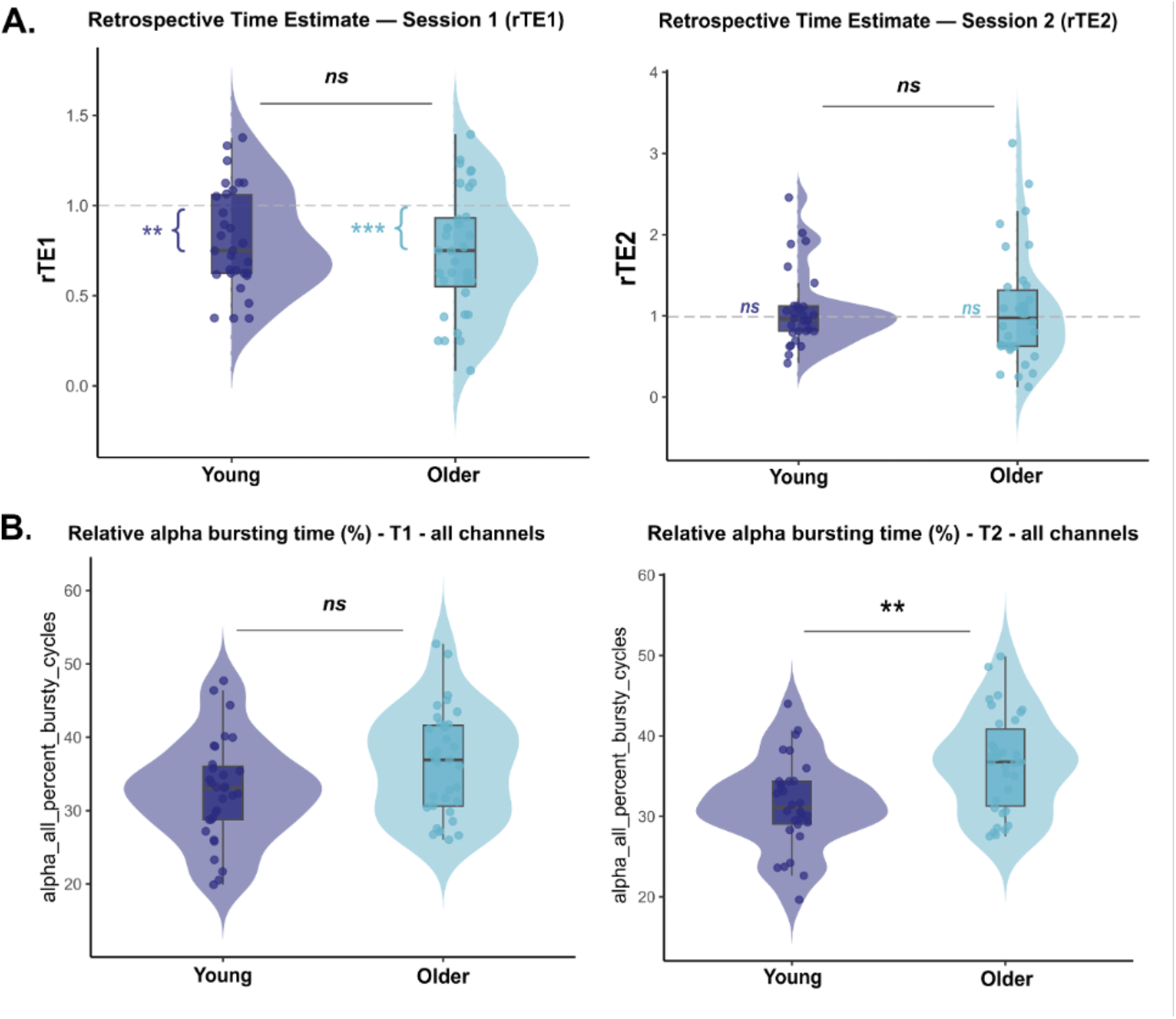
No behavioral difference for temporal perception but different resting-state EEG burst dynamics between younger and older adults. a) Violin plots showing the distribution of temporal measures across groups: No significant group difference for retrospective time estimates at Session 1 (rTE1, *p* = .666) and Session 2 (rTE2, *p* = .137). b) Violin plots showing the distribution of alpha burst dynamics across groups at T1 (left) and T2 (right): Robust group differences at T2 showing greater alpha burst activity in older compared to younger adults (*p* < .001, *p*FDR = .006, *d* = −0.88) and following the same direction but not surviving FDR correction at T1 (*p*FDR > .089). **Note**. Younger adults (n = 31, dark blue); Older adults (n = 36, light blue). Boxplots indicate the median (central line), interquartile range (box), and whiskers extending to 1.5 x the interquartile range, with individual dots representing participants. A ratio of 1.0 (dashed line) indicates accurate estimation; values below 1.0 indicate underestimation. Colored asterisks reflect one-sample tests against veridical duration (ratio = 1) within each group, whereas black indicators reflect between-group comparisons. Significance indicators reflect FDR-corrected *p*-values and are indicated as follows: *p* < .05*, *p* < .01**, *p* < .001***; ns = not significant.

The mixed ANCOVA on rTE (rTE1, rTE2) examining the potential expectation effects across sessions revealed a significant main effect of Session (*F*(1, 59) = 14.72, *p* < .001, ηp^2^ = .200), reflecting a general tendency to produce longer duration estimates during the retrospective duration at the second session relative to the first, regardless of group. Years of education significantly contributed to the model (*F*(1, 59) = 4.16, *p* = .046, ηp^2^ = .066). Neither the main effect of Group (*F*(1, 59) = 0.56, *p* = .459) nor the Group x Session interaction (*F*(1, 59) = 1.29, *p* = .260) reached significance. Post-hoc comparisons (Session 1 - Session 2) revealed that only older adults showed a significant increase in retrospective time estimates from Session 1 to Session 2 (*t(*59) = −3.31, *p* = .002), whereas no such change was observed in younger adults (*t*(59) = - 1.33, *p* = .187).

### 3.2. Naturalistic prospective encoding of time

#### Both groups underestimated naturalistic durations, with older adults showing relatively weaker underestimation for longer duration clips compared to younger adults

The mixed-design ANOVA (see Fig. 4) revealed no significant main effect of Group (*F*(1, 59) = 0.17, *p* = .682, ηp^2^ = .003), or Duration (*F*(2.27, 133.84) = 1.24, *p* = .296, ηp^2^ = .021). A significant main effect of Eventfulness was found (*F*(1, 59) = 25.79, *p* < .001, ηp^2^ = .304), with higher ratios for low-than high-eventfulness clips. Significant two-way interactions were observed for Group x Duration (*F*(2.27, 133.84) = 4.80, *p* = .007, ηp^2^ = .075), Group x Eventfulness (*F*(1, 59) = 4.56, *p* = .037, ηp^2^ = .072), and Duration x Eventfulness (*F*(2.72, 160.65) = 3.91, *p* = .012, ηp^2^ = .062). The three-way interaction was not significant (*p* = .232).

**Fig. 4.**
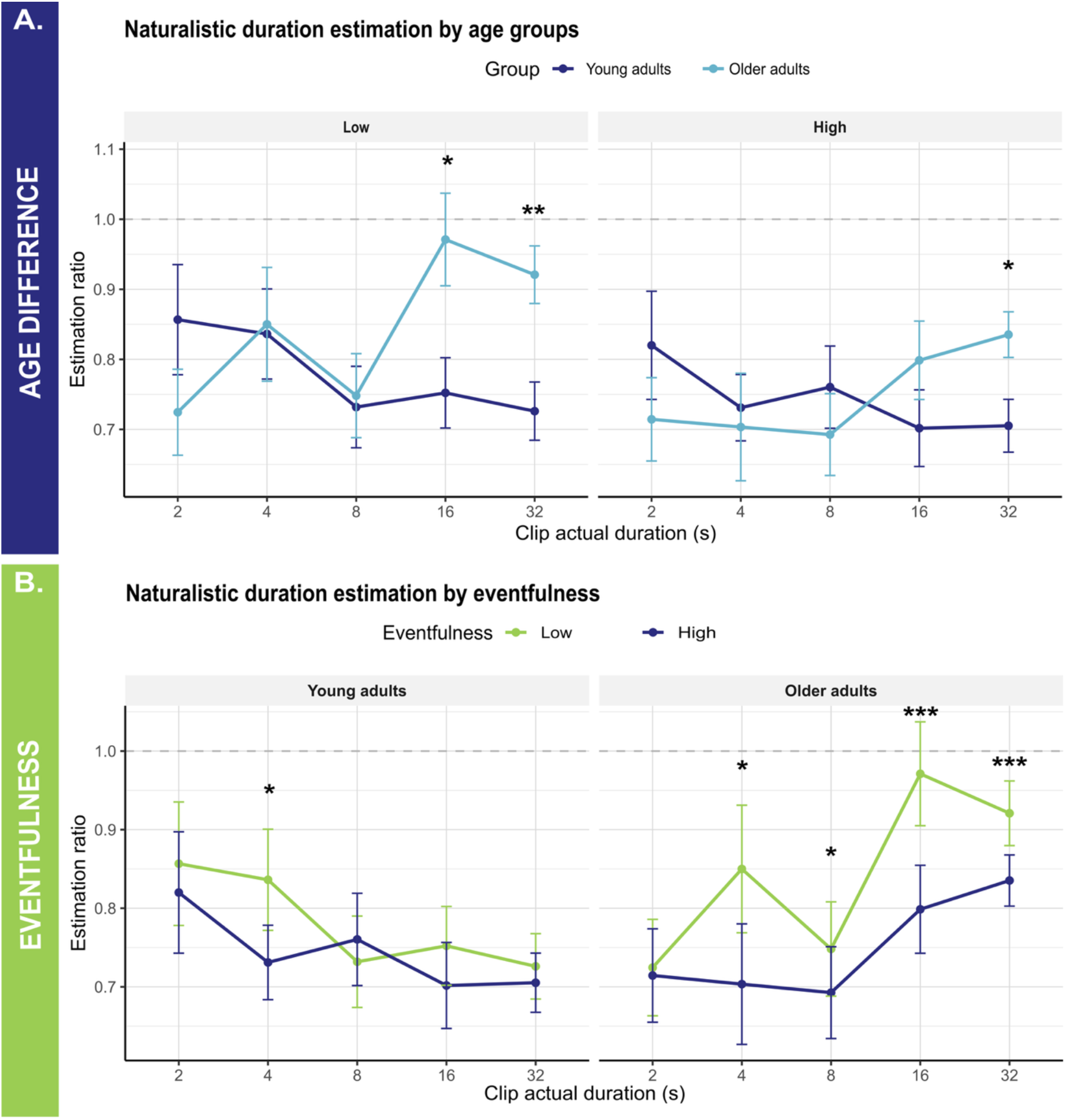
Naturalistic duration estimation by age groups and eventfulness conditions. a) *Age-group differences.* Line plots showing mean duration estimation ratios for the naturalistic video clips (estimated/actual duration, ± SE) as a function of clip duration (2-32 s), shown separately for low- and high-eventfulness clips, in younger and older adults. Older adults showed significantly higher estimations ratio than younger adults for long low-eventfulness clips (16-s: *p* = .042; 32-s: *p* = .002) and long high-eventfulness clips (32-s: *p* = .037). b) *Eventfulness differences.* Line plots showing mean duration estimation ratios for the naturalistic video clips (estimated/actual duration, ± SE) by eventfulness (low vs. high) and as a function of clip duration (2-32 s), shown separately for younger and older adults. Older adults showed significantly higher ratios for low-than high-eventfulness clips at 4-s (*p* = .032), 8-s (*p* = .043), 16-s (*p* < .001), and 32-s (*p* < .001), while younger adults showed this difference only at 4-s (*p* = .048). **Note.** Asterisks indicate significant post-hoc pairwise comparisons (Holm-corrected): \**p* < .05, \*\**p* < .01, \*\*\**p* < .001. Error bars represent ± 1 SE. A ratio of 1.0 (dashed line) indicates accurate estimation; values below 1.0 indicate underestimation. YA = younger adults (n = 31); OA = older adults (n = 36).

Post-hoc comparisons between groups revealed two main patterns. First, regarding group difference (Fig. 4a), older adults showed significantly higher ratios than younger adults for long low-eventfulness clips (16-s: *t*(59) = −2.08, *p* = .042; 32-s: *t*(59) = −3.21, *p* = .002) and long high-eventfulness clips (32-s: *t*(59) = −2.13, *p* = .037). Second, regarding eventfulness effects within each group (Fig. 4b), older adults showed significantly higher ratios for low-than for high-eventfulness clips across most durations (all *ps* < .043), whereas younger adults showed this difference only at 4-s clips (*t*(59) = 2.02, *p* = .048).

### 3.3. Oscillatory dynamics of retrospective encoding of time

#### No difference in alpha power or iAPF, but more alpha burst activity at rest for older than younger adults

Neither individual alpha peak frequency (iAPF) nor alpha power differed significantly between younger and older adults at T1 (iAPF: *t*(63) = 0.61, *p* = .545; alpha power: *t*(65) = 0.93, *p* = .355) or T2 (iAPF: *t*(61) = 0.66, *p* = .512; alpha power: *t*(61) = 0.48, *p* = .631), indicating that the two age groups in our sample did not differ in their baseline spectral characteristics.

Alpha burst dynamics were then compared between groups at T1 and T2 (Fig. 3b). Older adults showed higher percentage of alpha bursty cycles, principally at the whole-scalp level, compared to younger adults at T2 (all-channel: *p* < .001, *p*FDR = .006, *d* = −0.88; cluster-level: *p* = .027, *p*FDR = .074, *d* = −0.56). At T1, differences followed the same direction but did not survive FDR correction (all-channel: *p* = .032, *p*FDR = .089, *d* = −0.54; cluster-level: *p* = .111, *p*FDR = .233; *d* = −0.40). Complete descriptive statistics and group comparisons for EEG spectral measures and alpha burst dynamics are reported in Supplementary Table S3.

#### Alpha burst dynamics predicted retrospective time overestimation in older adults under genuine incidental encoding conditions

Partial correlations between temporal measures and session-matched resting-state EEG metrics revealed no significant associations for rTE1 or rTE2 in either group after controlling for years of education and expectancy (all *p*s > .13; Fig. 5a).

**Fig. 5.**
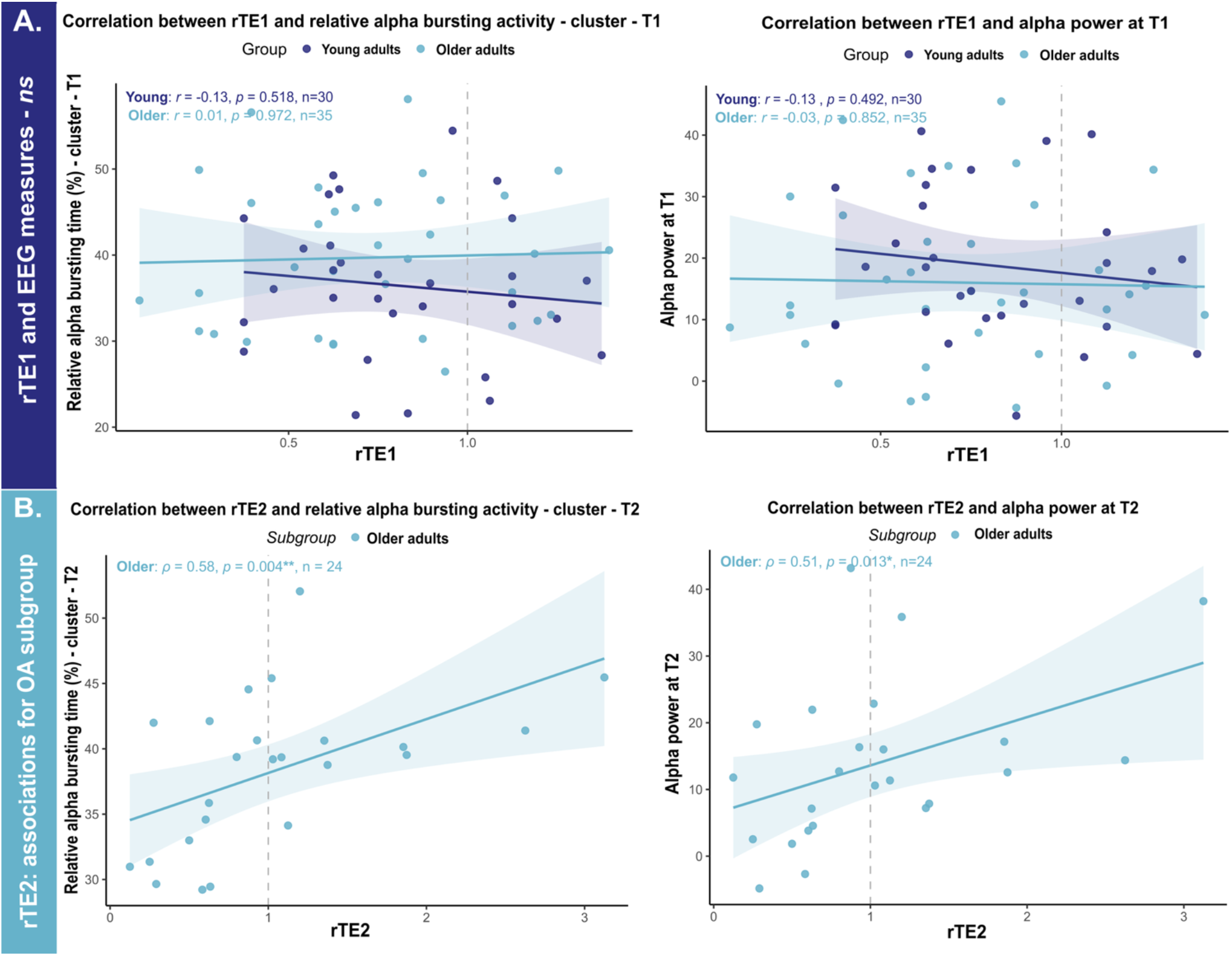
Associations between temporal measures and resting-state EEG metrics. a) Scatterplot with linear fit showing the absence of significant associations between rTE1 and alpha burst activity (left: relative alpha bursting time, cluster-level; right: alpha power) at T1 in both groups. b) In the subgroup of older adults who did not expect the timing question at T2 (n = 24), scatterplot with linear fit showing the robust positive associations between rTE2 and alpha burst dynamics at T2: relative alpha bursting time (*ρ* = .58, *p* = .004, *p*FDR = .010) and alpha power at T2 (*ρ* = .51, *p* = .013, *p*FDR = .038). **Note**. Reported *r, ρ* and *p* values correspond to partial correlations controlling for years of education (and expectancy for rTE2). Younger adults = dark blue; Older adults = light blue. All = all channels; Pool = cluster-level. Significance levels are indicated as follows: *p* < .05*, *p* < .01**, *p* < .001***.

Given the significant session effect observed in the mixed ANCOVA on rTE (*F*(1, 59) = 14.72, *p* < .001) and the context-dependent nature of retrospective time estimation [75], an exploratory analysis was conducted in the subgroup of participants who did not expect the timing question at Session 2. This subgroup consisted primarily of older adults (OA: n = 24, 67%; YA: n = 7, 23%). Due to the small younger adults’ subsample (n = 7), which prevented significant group comparisons, analyses were restricted to older adults (Fig. 5b). Partial correlations between rTE2 and EEG metrics at T2, controlling for years of education, revealed robust positive associations between rTE2 and multiple alpha burst metrics at T2 (all *p*FDR < .027), such as a higher percentage of alpha bursty cycles (*ρ* = .58, *p* = .004, *p*FDR = .010) is associated to overestimation of the video clips. Alpha power also showed a significant positive association with rTE2 (*ρ* = .51, *p* = .013, *p*FDR = .038) in this subgroup. Exploratory analyses examining whether subjective experience dimensions (boredom, arousal, richness) at T1 accounted for the absence of EEG-time associations are reported in Supplementary Table S4.

### 3.4. Oscillatory dynamics of naturalistic prospective encoding of time

#### Same levels of alpha burst activity for younger and older adults during naturalistic prospective duration estimation task, with alpha burst activity increasing with clip duration in both groups

No group difference was observed for alpha burst dynamics (*p* = .844). Interestingly, alpha burst activity increased with clip duration (*F*(3.30,184.94) = 47.23, *p* < .001, ηp^2^ = .458) and a significant Group x Duration interaction (*F*(3.30,184.94) = 3.54, *p* = .013, ηp^2^ = .059) was found, reflecting a greater increase in alpha bursts with clip duration in younger adults compared to older adults (Fig. 6a). However, no correlation between video estimation ratios and alpha burst activity reached significance in either group (all *p*s > .80 in younger adults; all *p*s > .95 in older adults).

**Fig. 6.**
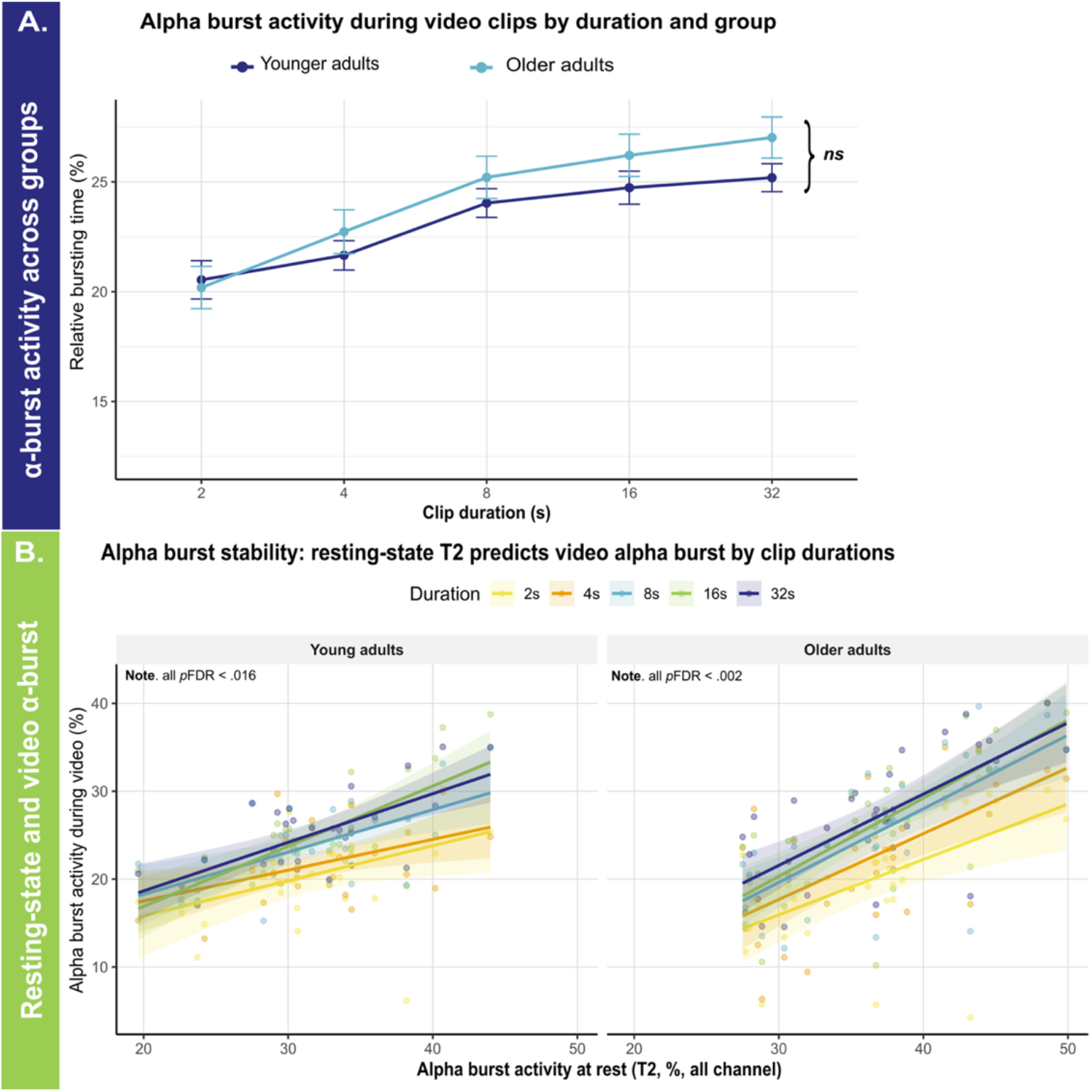
Oscillatory burst dynamics during the video-based duration estimation task. a) Line plots showing mean relative alpha bursting time (% bursty cycles ± SE) as a function of clip duration in younger and older adults. Alpha burst increase followed clip duration in both groups (*p* < .001) but was more gradual in older adults (Group x Duration: *p* = .013). b) Scatterplot with linear fit showing the relationship between resting-state alpha burst activity at T2 (all channels) and alpha burst activity during video clips, by duration and group. Resting-state alpha burst activity predicted alpha burst activity during the video-based duration estimation task across all durations in both groups (all *p*FDR < .016).

### 3.5. Associations between retrospective timing, naturalistic duration estimation, and neuropsychological measures

#### Resting-state alpha burst activity predicted alpha burst activity during the video-based duration estimation task in both groups

To explore whether individual alpha dynamics represent a stable disposition engaged across temporal contexts, we examined whether resting-state alpha burst activity was associated with alpha burst activity during naturalistic video encoding. This exploratory analysis directly addresses our broader aim of linking temporal reconstruction across incidental and naturalistic contexts.

In both groups, resting-state alpha burst activity at T2 was robustly and positively associated with alpha burst activity during the video-based duration estimation task across all clip durations (Fig. 6b). In younger adults, resting-state alpha burst activity (all channels) correlated significantly with alpha burst activity across durations (*r* and *ρ* ≥ .50, *p*FDR ≤ .016). In older adults, these associations were even stronger (*r* and *ρ* ≥ .57, *p*FDR ≤ .002). A comparable pattern was observed using the cluster-level alpha metrics in both groups (all *p*FDR ≤ .019).

#### Retrospective time overestimation at Session 2 predicted long naturalistic duration overestimation in older adults

To explore whether retrospective and naturalistic temporal reconstruction draw on shared processes, we examined associations between retrospective time estimates (rTE1, rTE2) and naturalistic duration ratios, focusing on long-duration clips where memory-based reconstruction is most engaged. This exploratory analysis directly addresses our aim of characterizing how temporal reconstruction operates across incidental and naturalistic contexts in aging.

In older adults, rTE2 was positively associated with long-duration ratios showing group differences (32-s high-eventfulness: *ρ* = .42, *p* = .017, *p*FDR = .047; 32-s low-eventfulness: *ρ* = .40, *p* = .024, *p*FDR = .047), suggesting that older adults who overestimated the retrospective duration of the session also overestimated long naturalistic clips (Fig. 7a,b). No significant associations were observed in younger adults (all *p*s > .436).

**Fig. 7.**
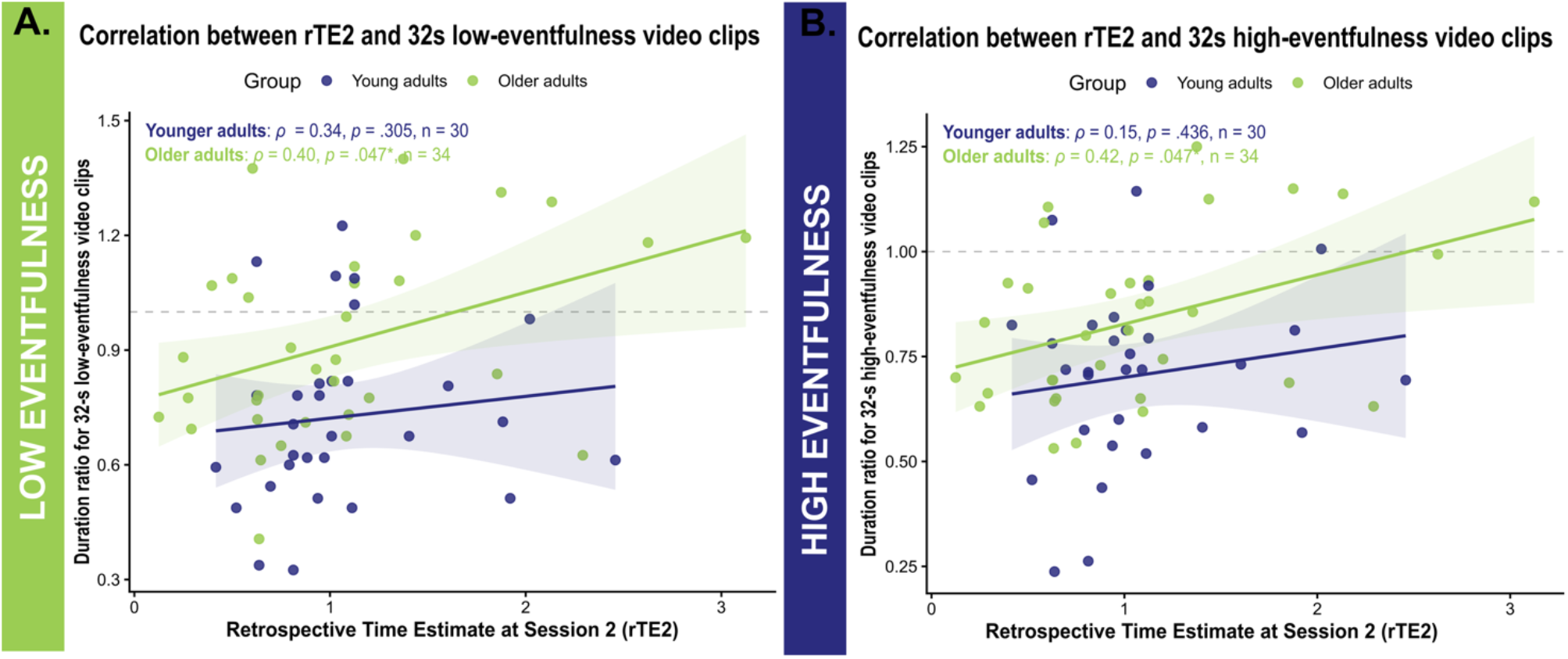
Partial correlations between retrospective time estimation at Session 2 (rTE2) and duration ratios for 32-second video clips, by eventfulness level and age group. Scatter plots showing the relationship between rTE2 and duration ratios for low-eventfulness a) and high-eventfulness b) video clips in younger (dark blue) and older adults (green). Regression lines are shown with 95% confidence intervals. The dashed horizontal line indicates accurate duration estimation (ratio = 1). **Note**. Reported *r, ρ* and *p* values correspond to partial correlations controlling for years of education (and expectancy for rTE2). *p*-values are FDR-corrected. Significance levels are indicated as follows: *p* < .05*, *p* < .01**, *p* < .001***.

Partial correlations between temporal measures (rTE1, rTE2) and neuropsychological indices revealed no significant associations after FDR correction in either group. For the video-based duration estimation task, significant associations with neuropsychological indices are reported in Supplementary Table S4.

#### Retrospective and naturalistic duration estimation share common variance in older adults, independently predicted by episodic memory

To further examine the relationship between retrospective and naturalistic duration estimation, we tested whether rTE2 predicted estimation of long video clips (32-s clips) in older adults, given the positive association observed between these measures. Two hierarchical regressions were conducted with the 32-s high- and low-eventfulness duration ratio as the dependent variable, years of education included as a covariate throughout (see Table 2).

**Table 2.**
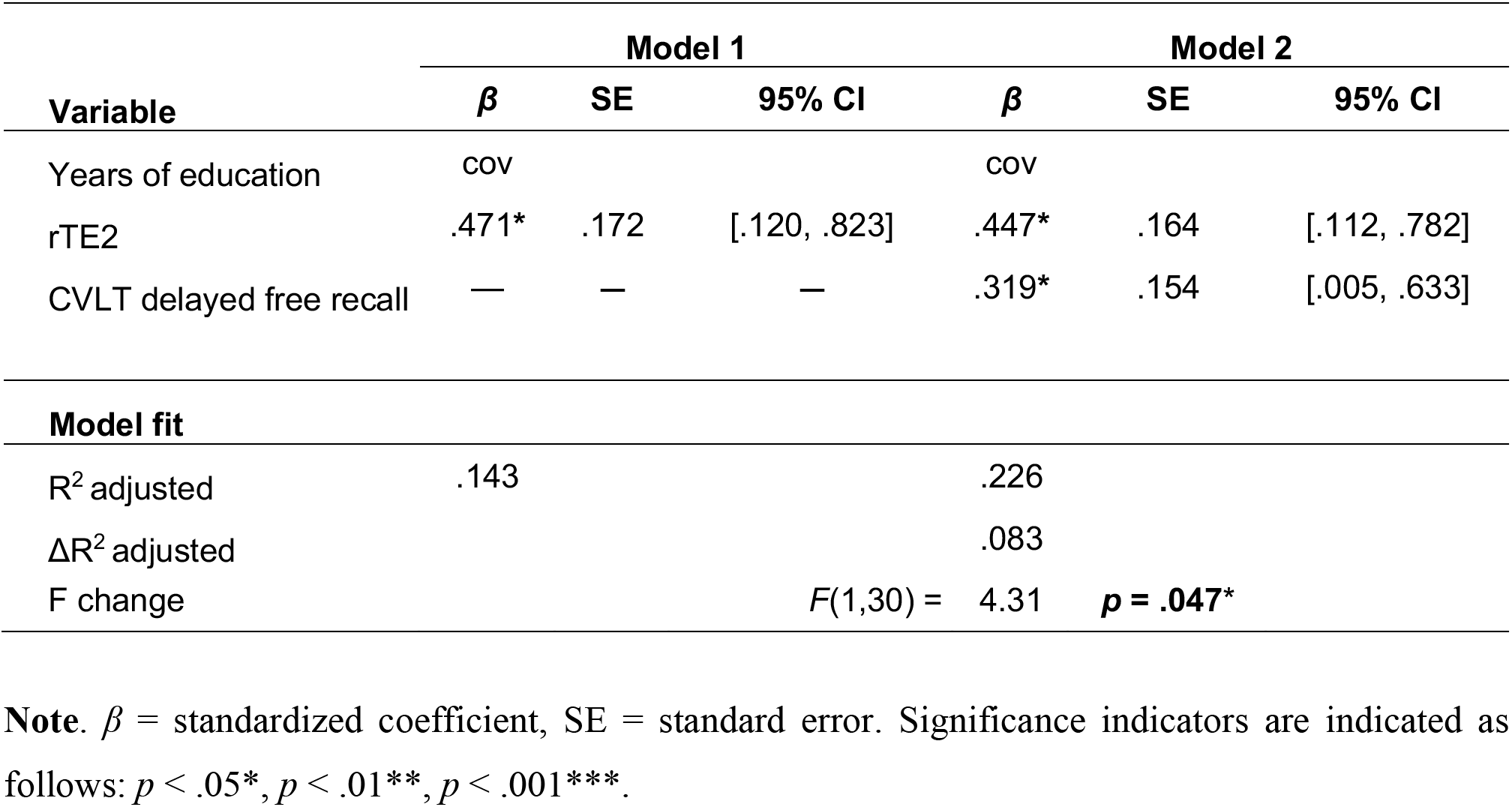
Hierarchical regression predicting naturalistic high-eventfulness duration estimation from retrospective timing and episodic memory in older adults. Results of a two-step hierarchical regression with the 32-s high-eventfulness duration ratio as the dependent variable and years of education included as a covariate throughout. Step 1 included rTE2 as predictor; Step 2 additionally included delayed episodic memory (CVLT delayed free recall).

For the 32-s low-eventfulness condition, rTE2 emerged as a significant predictor in Step 1 (*β* = .395, *p* = .031, Δ*R^2^* adjusted = .111). However, the addition of delayed episodic memory in Step 2 did not significantly improve model fit (Δ*R^2^*adjusted = .025, *p* = .178, *F*(1,30) = 1.90, *p* = .178).

For the 32-s high-eventfulness condition, rTE2 significantly predicted the 32-s high-eventfulness duration ratio in Step 1 (*β* = .471, *t*(31) = 2.74, *p* = .010, *R*^2^ adjusted = .143). In Step 2, adding delayed episodic memory score (CVLT delayed free recall) significantly improved model fit (ΔR^2^ adjusted = .083, *F*(1,30) = 4.31, *p* = .047), with both rTE2 (*β* = .447, *p* = .010) and CVLT (*β* = .319, *p* = .047) remaining significant independent predictors (total *R*^2^ adjusted = .226), suggesting that overestimation of long high-eventfulness naturalistic durations in older adults is influenced by both their general retrospective duration judgements abilities and their episodic memory resources. This pattern held after controlling for years of education.

In summary, retrospective time estimates (rTE1, rTE2) did not differ between age groups. For naturalistic durations, older adults showed higher estimation ratios than younger adults for longer video clips (16-s and 32-s), particularly under low eventfulness condition. Resting-state alpha burst activity was higher in older adults, and both alpha burst activity and alpha power predicted retrospective time estimates in this group. Resting-state alpha burst activity predicted alpha burst activity during video encoding in both groups. Finally, retrospective and naturalistic duration estimates were positively associated in older adults, with episodic memory contributing additional predictive variance for long high-eventfulness clips (32-s).

## 4. Discussion

The present study aimed to investigate how retrospective and naturalistic temporal encoding are shaped by advancing age, and whether episodic memory resources constitute a common substrate linking these two forms of temporal reconstruction. To this end, we examined retrospective time estimation and naturalistic duration estimation in younger and older adults, alongside their resting-state and task-related oscillatory correlates. Behaviorally, we anticipated limited mean-level age differences but greater inter-individual variability, with oscillatory dynamics differing across age groups and predicting duration estimation. We also hypothesized that episodic memory decline would selectively impact the most memory-dependent conditions and influence temporal reconstruction. Overall, our results partially supported these working hypotheses. While younger and older adults reconstructed short intervals similarly, aging selectively shifted naturalistic duration estimation, with older adults producing longer duration estimates for longer video clips compared to younger adults, especially for low-eventfulness clips. At the neural level, resting-state alpha burst dynamics differed between age groups, and greater alpha burst activity in older adults, as well as higher alpha power, were predictive of retrospective time estimation in older adults. Additionally, resting-state burst activity was associated with alpha burst activity during video clips in both groups. Finally, associations between retrospective and naturalistic duration estimation, further modulated by episodic memory resources, were observed in older adults, suggesting that temporal reconstruction across timescales draws on a common age-sensitive substrate.

At the behavioral level, both groups showed similar retrospective time estimates with a general tendency to underestimate the duration, as expected when one is not paying attention to time [15,113]. Older adults showed similar mean and variability in their estimations compared to younger adults, consistent with results on retrospective timing in aging showing little to no systematic age difference [19,37,53,114]. On the other hand, contrary to our working hypothesis, they did not show greater inter-individual variability. This result may reflect the cognitively healthy profiles of our older adult sample (MoCA ≥ 26), as well as the incidental nature of the task, which limits the influence of explicit timing strategies and attentional demands that typically introduce variability in prospective paradigms [54]. Retrospective time estimates nonetheless increased from Session 1 to Session 2, driven by older adults, likely reflecting the accumulation of contextual changes across the experimental session rather than improved temporal tracking [6,115,116]. Indeed, richer contextual encoding during the second visit, including familiarity with the lab environment, as well as less stress-related performance, prior experience with the task, and accumulated experimental landmarks, may have provided additional retrieval cues supporting longer duration estimates. Importantly, most older adults did not anticipate the timing question at Session 2, suggesting that their estimates reflected genuine retrospective reconstruction, aligning with the view that under truly incidental encoding conditions, retrospective time estimation captures the richness of the encoded episode more faithfully [75].

With regards to naturalistic durations, both groups underestimated the length of video clips, with older adults showing relatively longer estimates than younger adults for longer duration clips, particularly under low eventfulness. This pattern contrasts with studies attributing age-related timing differences to attentional overload when the context is rich and attentional control is taxed [36,37,60] and instead suggests that the critical factor here is the reduced availability of memorable events to support prospective duration reconstruction, with older adults showing greater sensitivity to their absence under low-eventfulness conditions. Importantly, age-related differences were more pronounced for longer durations, that is when temporal reconstruction relies more heavily on memory-based mechanisms [1,2,23]. As clip length increases beyond working memory span, duration judgments must be reconstructed from episodic traces that are themselves temporally compressed in memory [38,117]. The influence of eventfulness further supports this interpretation: older adults showed reduced underestimation for low-than high-eventfulness clips across most durations, suggesting that when fewer events are available to structure the encoded episode [40,41] temporal reconstruction is particularly challenged in older adults compared to younger adults. Notably, this pattern runs counter to predictions based on episodic compression accounts, which would expect reduced episodic resources in older adults to produce greater underestimation [38,117], suggesting that alternative mechanisms may contribute to naturalistic duration estimation in aging. Rather, this pattern is consistent with an increased reliance on external environmental support and cues to keep track of time, in line with the environmental dependency account of cognitive aging [118]. Together, these findings suggest that naturalistic duration estimation in older adults is particularly sensitive to conditions where memory-based reconstruction is most demanded, namely long durations and impoverished event structure, even if the precise mechanisms underlying this sensitivity remain to be fully elucidated.

At the neural level, we found no difference in alpha power or iAPF at rest between younger and older adults, but different resting-state and task-related burst dynamics across age groups. The absence of group differences in iAPF and alpha power contrasts with previous research demonstrating age-related slowing of alpha peak frequency and alterations in alpha power [119,120], which may reflect the relatively young age of our older adult sample (mean age 69 years, range 61-79). Notably, such effects have been shown to follow non-linear trajectories becoming more pronounced in older age ranges [121]; a pattern consistent with studies reporting robust iAPF slowing in samples spanning broader age ranges, often extending into the eighth and nineth decades of life [69,122,123]. Additionally, by isolating the periodic component prior to estimation of iAPF and alpha power using the IRASA algorithm, our approach used a more conservative but methodologically cleaner estimate of alpha dynamics, as aperiodic changes with age can otherwise influence broadband power and apparent peak frequency [111,124] and could explain the absence of replication on our side.

Beyond spectral power, the temporal structure of oscillatory activity (i.e., whether rhythms occur in sustained bursts or isolated cycles) provides additional information about neural dynamics that traditional Fourier-based approaches could not capture [71,111]. Using a cycle-by-cycle burst analysis, we found that older adults showed greater resting-state alpha burst activity compared to younger adults, and similar alpha levels during the naturalistic duration estimation task. Our findings extend previous work on age-related spectral changes by showing that burst dynamics dissociated across age groups in ways that are not captured by power alone, consistent with the view that oscillatory bursts reflect functionally distinct neural states [71,125]. Resting-state neural timescales have been shown to constrain how durations are integrated and segmented during perception and cognition, with intrinsic temporal dynamics at rest persisting into task states [126], suggesting that individual differences in resting-state burst activity may shape temporal processing more broadly.

Unlike the video-based duration estimation task, the retrospective timing paradigm offers no opportunity for active maintenance, as participants are unaware that a duration judgment will be required. In this context, we found that resting-state alpha burst activity and alpha power predicted longer retrospective time estimates in older adults under genuine incidental encoding conditions. The absence of such associations in the full sample relates to the sensitivity of alpha burst metrics to contextual factors modulating the quality of incidental encoding [75], as discussed previously. The alpha estimation associations restricted to genuine incidental conditions in older adults in this study suggests that oscillatory markers of temporal reconstruction may be particularly sensitive to aging under conditions of minimal attentional engagement, complementing behavioral findings showing that explicit prospective timing tasks often mask age-related differences that emerge under more naturalistic or memory-dependent conditions [56,57]. Importantly, the predictive role of alpha burst activity in retrospective time estimation aligns with the episodic stamping hypothesis, according to which spontaneous alpha bursts index the neural mechanism by which temporal information is passively tagged onto episodic memory traces during ongoing experience [74,75]. Consistent with this view, alpha burst activity increased with clip duration during video viewing in both groups and resting-state alpha burst activity robustly predicted alpha burst activity during the video task across all durations, corroborating that individual alpha dynamics constitute a stable neural disposition shaping temporal processing across contexts [16,127]. This increased alpha engagement may further reflect a tendency toward internally directed cognition at rest, consistent with evidence linking alpha oscillations to mind-wandering states [128,129], which could in turn facilitate incidental temporal encoding. In line with Azizi and collaborators [74], these findings suggest that alpha burst activity may constitute a memory-based time-tracking mechanism engaged not only at rest but also during events that exceed working memory capacity, and that this mechanism operates across timescales of temporal reconstruction.

Associations between retrospective and naturalistic duration estimation confirmed that older adults who overestimated the retrospective duration of the session also overestimated long naturalistic clips, suggesting that these two forms of temporal reconstruction share a common substrate in this age group. Hierarchical regression analyses revealed that this shared variance was further modulated by episodic memory resources. In particular, for 32-s high-eventfulness clips, delayed episodic memory scores contributed significant independent predictive variance beyond retrospective timing, with both predictors remaining significant in the final model. This pattern suggests that the reconstruction of long naturalistic durations in older adults draws jointly on their general tendency to reconstruct time retrospectively and on their episodic memory resources. This result supports the hypothesis that temporal reconstruction becomes increasingly memory-dependent as duration increases, providing empirical support for recent theoretical accounts proposing that the brain continuously stamps temporal information onto episodic representations during ongoing experience [2,13]: when episodic resources are sufficiently preserved, they contribute independently to the reconstruction of long naturalistic durations, whereas their age-related decline [58,63] may reduce the fidelity of this temporal stamping process. Furthermore, the absence of an independent episodic memory contribution for low-eventfulness clips complements this idea with frameworks suggesting that impoverished event structure limits the availability of episodic anchors on which memory-based reconstruction can operate [8,33,37], such that even intact episodic resources cannot compensate for the lack of temporal scaffolding. Under the contextual change model, these events are considered to serve as contextual anchors for duration reconstruction [116,130], and their reduced availability under low eventfulness may disproportionately affect older adults whose episodic memory resources are more limited. Interestingly, while recent theoretical accounts predict that reduced experiential density with aging should increase temporal compression, whether at the scale of life episodes [131] or of individual events [38], our older adults showed the opposite pattern, underestimating long naturalistic durations less than younger adults, highlighting the multifactorial nature of naturalistic duration estimation in aging. Furthermore, recent work suggests that temporal compression in episodic memory cannot be fully accounted for by central tendency effects alone, pointing instead to nested contextual changes as a key driver [117]; a framework that may help shed light on the shifted naturalistic estimation pattern observed in our older adults. In line with this, older adults have been shown to rely less on temporal structure when recalling naturalistic events [132], potentially weakening the temporal scaffolding on which duration reconstruction depends. Taken together, these findings converge on a coherent picture in which retrospective timing and naturalistic duration estimation draw on overlapping reconstructive processes, with episodic memory serving as a ke moderator of temporal judgment accuracy in older adults, particularly when the richness of the encoded experience provides sufficient structure for memory-based reconstruction to operate.

### Limitations

While this study offers a novel characterization of how retrospective and naturalistic temporal encoding are reshaped with advancing age at both behavioral and oscillatory levels, some limitations should be acknowledged. First, the high proportion of younger adults who anticipated the timing question at Session 2 resulted in a subgroup too small to conduct equivalent analyses under genuine incidental conditions (n = 7), preventing direct comparison with older adults and limiting the generalizability of the alpha burst-retrospective timing association to this age group. Future studies will be needed to further characterize retrospective timing across repeated sessions in younger adults. Relatedly, as previously noted, older adults may have been more familiar with the experimental procedure at the second session, which could partly account for the stronger associations observed at T2 compared to T1. However, disentangling session effects from increased procedural familiarity is inherently constrained by the retrospective nature of the paradigm, which cannot be fully replicated once participants are no longer naive to the timing question. Second, although the subgroup of older adults who did not anticipate the timing question represents the ideal conditions for capturing memory-based temporal reconstruction, this subgroup was identified post-hoc, which introduces a potential selection bias. However, the fact that this subgroup comprised the majority of our older adults (n = 24, 67%) and that their non-anticipation was verified experimentally attenuates this concern. Third, our older adult sample was relatively young (mean age 69 years, range 61-79), which may have limited the expression of age-related oscillatory changes that typically become more pronounced in older age ranges. Extending this paradigm to oldest-old participants would allow a more comprehensive characterization of how temporal reconstruction and its neural substrates are reorganized across the full aging spectrum. Finally, neuropsychological assessments revealed no direct correlations between episodic memory scores and temporal task performance, likely reflecting the ceiling effects in our groups inherent to a cognitively healthy sample. The contribution of episodic memory resources to each form of temporal reconstruction remains to be investigated in populations with more marked cognitive decline and greater variability in educational background, where greater variability in memory performance may unmask differential associations with retrospective and naturalistic duration estimation.

### Conclusion

By combining retrospective timing and naturalistic duration estimation, alongside resting-state and task-related oscillatory measures, this study addresses a critical gap in the literature on how retrospective and naturalistic temporal encoding differ in their sensitivity to advancing age. The present findings cast individual alpha burst dynamics as a stable neural disposition functioning as a memory-based time-tracking mechanism, engaged both incidentally at rest and prospectively during naturalistic encoding, with aging selectively unmasking its role under memory-demanding conditions. Our results demonstrate that while behavioral performance was largely preserved in older adults, age-related reorganization manifested at the oscillatory level in ways that behavioral measures alone could not capture. Importantly, we show that retrospective and naturalistic temporal reconstruction were found to draw on common episodic memory resources in older adults, with episodic memory contributing independent predictive variance to long naturalistic duration estimation beyond the shared reconstructive tendency. Future work combining multimodal neuroimaging will be needed to directly identify the neural generators of the observed burst dynamics and to determine the extent to which hippocampal, whose contributions are only indirectly reflected in surface EEG, and cortical networks support the temporal stamping mechanism proposed here, with potential implications for the early detection of age-related reorganization in temporal cognition.

## 5. Sources of funding

This work was supported by the French National Research Agency (ANR) as part of the PRC ULIS project (ANR-22-CE28-0029).

## 6. CRediT authorship contribution statement

**Conceptualization**: Florentine Fricker, Giulia Buzi, Raphaël Bordas, Nathan Leroy, Patrice Clochon, Arnaud D’Argembeau, Virginie van Wassenhove, Thomas Hinault; **Methodology**: Florentine Fricker, Thomas Hinault; **Software**: Raphaël Bordas; **Validation**: Raphaël Bordas, Thomas Hinault; **Formal analysis**: Florentine Fricker; **Investigation**: Florentine Fricker, Giulia Buzi, Maëlys Morantin, Franck Doidy; **Resources**: Patrice Clochon; **Data curation**: Florentine Fricker, Giulia Buzi; **Writing - original draft**: Florentine Fricker, Thomas Hinault; **Writing - review and editing**: Florentine Fricker, Giulia Buzi, Maëlys Morantin, Franck Doidy, Patrice Clochon, Raphaël Bordas, Nathan Leroy, Arnaud D’Argembeau, Virginie van Wassenhove, Thomas Hinault; **Visualization**: Florentine Fricker; **Supervision**: Arnaud D’Argembeau, Virginie van Wassenhove, Thomas Hinault; **Funding acquisition:** Thomas Hinault.

## 7. Declaration of Competing Interest

None.

## Footnotes

1 Neural bursts refer to transient, high-amplitude episodes of oscillatory activity that occur within an otherwise irregular signal, as opposed to sustained rhythmic activity. Such transient dynamics, detectable across multiple frequency bands including alpha, have been linked to a broad range of cognitive processes across development and aging, and their detection through cycle-by-cycle approaches captures aspects of neural activity that traditional spectral methods tend to obscure by averaging across time [71–73].

## Notes

### Competing Interest Statement

The authors have declared no competing interest.

