## Supplementary file for "Aging effects on retrospective and naturalistic encoding of time and their relationship with memory"

### Statistical analyses

**Figure S1. Subjective experience ratings and group comparisons during the retrospective timing task at Session 1.**

Left panel: Distribution of z-scored values for each experience dimension, presented as horizontal boxplots (median, interquartile range, individual data points) for younger (dark blue, n = 30-31) and older adults (light blue, n = 34-36). Scores were standardized (z-scored) across the full sample prior to plotting to allow visual comparison across dimensions with different scales. Right panel: Summary statistics and group comparison results.

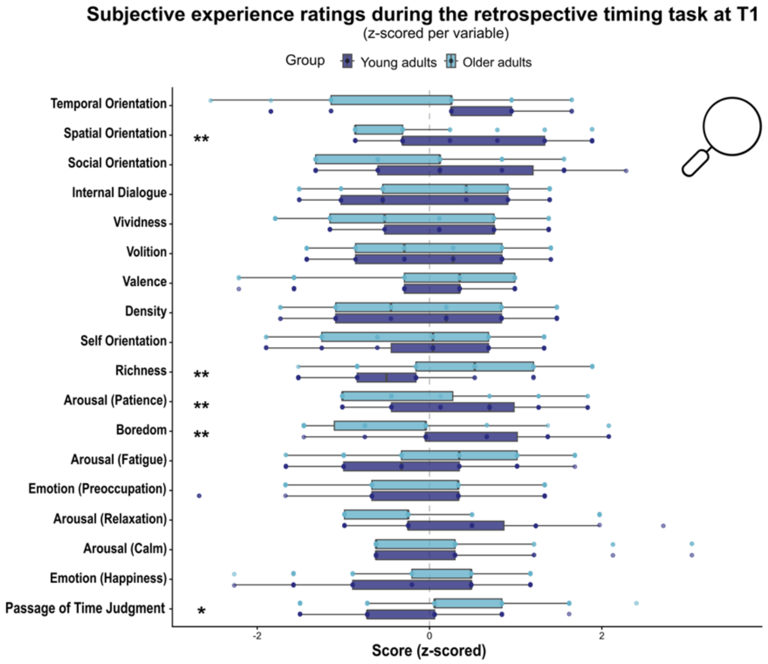


**Group comparisons for subjective experience ratings during the retrospective timing task at T1**

| Dimension | M (SD) YA | M (SD) OA | *W* | *r* | *p* | pFDR |
|---|---|---|---|---|---|---|
| **Richness** | **-0.476 (0.835)** | **0.408 (0.957)** | **265.0** | **-.50** | **< .001*** ** | **.006** ** |
| **Boredom** | **0.414 (0.905)** | **-0.366 (0.946)** | **796.5** | **.47** | **< .001*** ** | **.006** ** |
| **Spatial orientation** | **0.38 (1.052)** | **-0.327 (0.836)** | **803.0** | **.44** | **.001** ** | **.006** ** |
| **Arousal (Patience)** | **0.385 (0.916)** | **-0.331 (0.961)** | **805.0** | **.44** | **.001** ** | **.006** ** |
| **Passage of Time Judgment** | **-0.369 (0.802)** | **0.327 (1.053)** | **324.0** | **-.40** | **.004** ** | **.015* ** |
| Vividness | 0.318 (0.888) | -0.274 (1.021) | 743.0 | .33 | .018* | .054 |
| Arousal (Relaxation) | 0.207 (0.968) | -0.183 (1.005) | 688.5 | .27 | .052· | .134 |
| Arousal (Fatigue) | -0.239 (0.976) | 0.211 (0.987) | 403.5 | -.26 | .069· | .156 |
| Internal dialogue | -0.246 (0.998) | 0.218 (0.964) | 408.0 | -.25 | .080· | .160 |
| Social orientation | 0.213 (1.081) | -0.189 (0.896) | 653.0 | .20 | .145 | .252 |
| Valence | -0.126 (0.95) | 0.111 (1.043) | 435.0 | -.20 | .154 | .252 |
| Density | 0.174 (1.054) | -0.154 (0.938) | 645.0 | .19 | .182 | .255 |
| Temporal orientation | 0.141 (1.022) | -0.125 (0.978) | 640.5 | .18 | .184 | .255 |
| Arousal (Calm) | -0.101 (0.922) | 0.087 (1.068) | 473.0 | -.10 | .435 | .560 |
| Self orientation | 0.083 (1.001) | -0.071 (1.008) | 568.5 | .08 | .563 | .675 |
| Emotion (Preoccupation) | -0.044 (1.063) | 0.039 (0.955) | 499.5 | -.02 | .890 | .964 |
| Emotion (Sadness) | -0.022 (1.053) | 0.019 (0.966) | 546.5 | .01 | .962 | .964 |
| Volition | 0.019 (0.98) | -0.017 (1.032) | 546.5 | .01 | .964 | .964 |

**Note.** M = mean; SD = standard deviation; *W* = Wilcoxon rank-sum statistic; *r* = rank-biserial correlation coefficient. *p*FDR = Benjamini-Hochberg corrected p-value. Significance levels are indicated as follows: $p < .05$*, $p < .01$**, $p < .001$***.

**Figure S2. Features of video excerpts used in the video-based duration estimation task.**

Left panel: Table of all 50 video excerpts included in the main experiment, characterized by their duration (2, 4, 8, 16, or 32 seconds), video type (*Binder*, *Breakfast*, *Se*, *Sweeping*, *Tire*), eventfulness condition (High, Low), and mean density of perceived event boundaries.

Right panel: Mean density of perceived event boundaries plotted as a function of excerpt duration, separately for each video type and eventfulness condition. Values reflect the average number of participants from Kurby & Zacks (2011) who marked an event boundary per second within each excerpt. High-eventfulness excerpts consistently show greater boundary density than low-eventfulness excerpts across durations and video types (mean boundary density across all excerpts: high = 6 participants/s; low = 1 participant/s).

| Duration (s) | Video type | Eventfulness | Mean density of perceived event boundaries |
|---|---|---|---|
| 32 | Binder | High | 3.88 |
| 32 | Binder | Low | 1.78 |
| 32 | Breakfast | High | 9.38 |
| 32 | Breakfast | Low | 3.94 |
| 32 | Se | High | 7.25 |
| 32 | Se | Low | 0.94 |
| 32 | Sweeping | High | 3.94 |
| 32 | Sweeping | Low | 0.31 |
| 32 | Tire | High | 4.00 |
| 32 | Tire | Low | 0.06 |
| 16 | Binder | High | 3.81 |
| 16 | Binder | Low | 1.75 |
| 16 | Breakfast | High | 9.06 |
| 16 | Breakfast | Low | 3.94 |
| 16 | Se | High | 7.63 |
| 16 | Se | Low | 1.06 |
| 16 | Sweeping | High | 3.94 |
| 16 | Sweeping | Low | 0.31 |
| 16 | Tire | High | 4.06 |
| 16 | Tire | Low | 0.31 |
| 8 | Binder | High | 3.88 |
| 8 | Binder | Low | 1.75 |
| 8 | Breakfast | High | 9.38 |
| 8 | Breakfast | Low | 3.25 |
| 8 | Se | High | 7.5 |
| 8 | Se | Low | 0.88 |
| 8 | Sweeping | High | 3.88 |
| 8 | Sweeping | Low | 0.25 |
| 8 | Tire | High | 4.00 |
| 8 | Tire | Low | 0.25 |
| 4 | Binder | High | 3.75 |
| 4 | Binder | Low | 1.50 |
| 4 | Breakfast | High | 9.50 |
| 4 | Breakfast | Low | 3.00 |
| 4 | Se | High | 7.25 |
| 4 | Se | Low | 0.25 |
| 4 | Sweeping | High | 4.00 |
| 4 | Sweeping | Low | 0.00 |
| 4 | Tire | High | 4.50 |
| 4 | Tire | Low | 0.00 |
| 2 | Binder | High | 4.00 |
| 2 | Binder | Low | 2.00 |
| 2 | Breakfast | High | 9.00 |
| 2 | Breakfast | Low | 3.00 |
| 2 | Se | High | 7.00 |
| 2 | Se | Low | 0.50 |
| 2 | Sweeping | High | 4.00 |
| 2 | Sweeping | Low | 0.00 |
| 2 | Tire | High | 4.00 |
| 2 | Tire | Low | 0.00 |

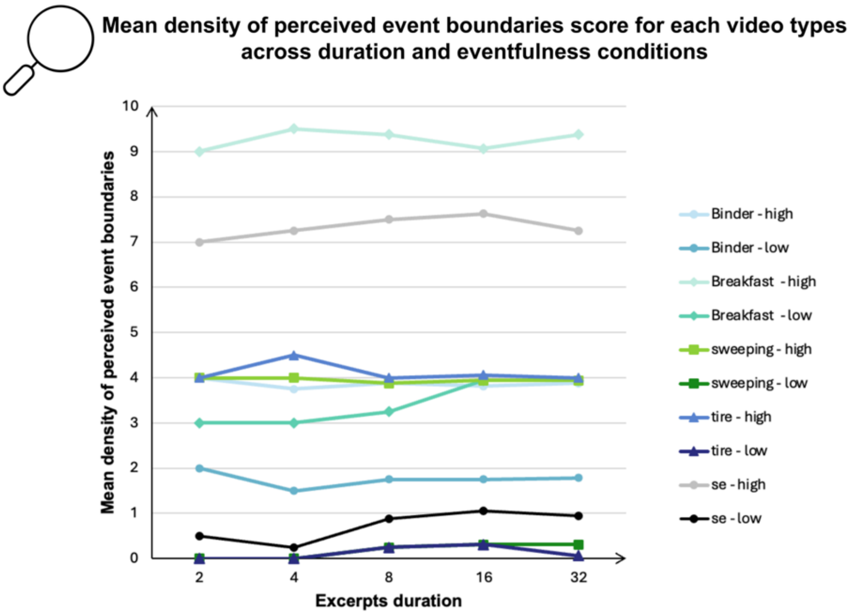

**Figure S3. Subjective experience ratings and group comparisons during the video-based duration estimation task at Session 2.**

Left panel: Distribution of scores for each experience dimension, presented as boxplots (median, interquartile range, individual data points) for younger (dark blue, n = 30) and older adults (light blue, n = 35). Right panel: Summary statistics and group comparison results.

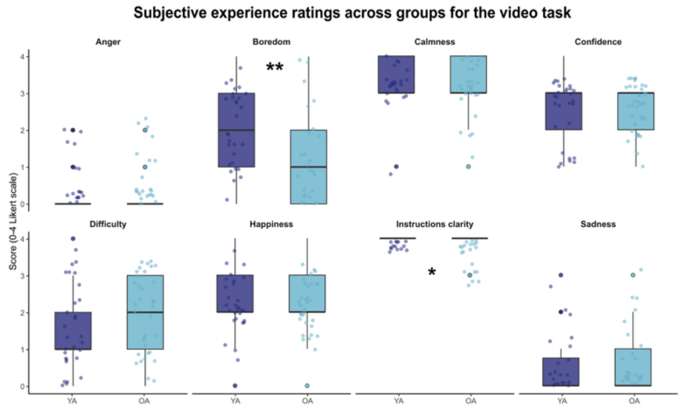


**Group comparison for subjective experience ratings during the video-based duration estimation task at Session 2**

| Variable | M (SD) YA | M (SD) OA | *W* | *r* | *p* | *p*FDR |
|---|---|---|---|---|---|---|
| Anger | 0.33 (0.71) | 0.34 (0.68) | 514.0 | r=-.02 | .847 | .968 |
| **Boredom** | 2.20 (1.03) | 1.11 (1.41) | 782.0 | r= .49 | **< .001*** ** | **.004** ** |
| Calmness | 3.30 (0.65) | 3.26 (0.78) | 528.0 | r= .01 | .971 | .971 |
| Confidence | 2.53 (0.94) | 2.57 (0.61) | 548.0 | r= .04 | .730 | .968 |
| Difficulty | 1.50 (1.14) | 1.86 (1.03) | 425.0 | r=-.19 | .172 | .459 |
| Happiness | 2.23 (0.86) | 2.17 (0.89) | 544.5 | r= .04 | .787 | .968 |
| **Instructions clarity** | 4.00 (0.00) | 3.80 (0.41) | 630.0 | r= .20 | **.010*** | **.042*** |
| Sadness | 0.40 (0.77) | 0.46 (0.82) | 511.0 | r=-.03 | .821 | .968 |

**Note.** M = mean; SD = standard deviation; *W* = Wilcoxon rank-sum statistic; *r* = rank-biserial correlation coefficient. *p*FDR = Benjamini-Hochberg corrected *p*-value. Significance levels are indicated as follows: $p < .05$*, $p < .01$**, $p < .001$***.

**Table S1. Test-retest reliability of burst-derived EEG metrics between Session 1 and Session 2, assessed using Intraclass Correlation Coefficients.**

Burst-derived metrics were computed separately for alpha band, across all channels and cluster-level electrodes. Overall, alpha burst quantity and duration metrics demonstrated good to excellent test-retest reliability in both age groups (ICC ≥ .75). In contrast, waveform symmetry indices (Burst PT symmetry and Burst RD symmetry) showed poor to low reliability (ICC < .40), especially in younger adults, suggesting that these metrics reflect state-dependent rather than trait-level oscillatory dynamics. ICC values were interpreted following standard benchmarks (Cicchetti, 1994): ICC < .40 = poor; .40-.59 = moderate; .60-.74 = good; ≥ .75 = excellent. CI = 95% confidence interval. Observations exceeding ±3 standard deviations from the group mean were treated as outliers and replaced with NA before test-retest reliability computation.

| Older Adults (OA) | | | | Younger Adults (YA) | | | |
|---|---|---|---|---|---|---|---|
| Alpha burst metrics | n | ICC | Label | Alpha burst metrics | n | ICC | Label |
| Percentage of burst samples (pool) | 34 | .915 [.837, .956] | *** (excellent) | Burst period (pool) | 30 | .870 [.748, .936] | *** (excellent) |
| Relative burst time (pool) | 34 | .913 [.834, .955] | *** (excellent) | Burst period (all) | 30 | .861 [.731, .931] | *** (excellent) |
| Burst period (all) | 34 | .893 [.798, .945] | *** (excellent) | Percentage of burst samples (all) | 30 | .810 [.641, .904] | *** (excellent) |
| Percentage of burst samples (all) | 34 | .885 [.784, .941] | *** (excellent) | Relative burst time (all) | 30 | .803 [.629, .901] | *** (excellent) |
| Bursts per second (pool) | 34 | .884 [.781, .940] | *** (excellent) | Percentage of burst samples (pool) | 30 | .783 [.595, .890] | *** (excellent) |
| Relative burst time (all) | 34 | .883 [.779, .939] | *** (excellent) | Relative burst time (pool) | 30 | .775 [.582, .886] | *** (excellent) |
| Bursts per second (all) | 34 | .881 [.776, .938] | *** (excellent) | Burst duration mean (all) | 29 | .773 [.575, .886] | *** (excellent) |
| Burst period (pool) | 34 | .878 [.772, .937] | *** (excellent) | Number of burst cycles (all) | 30 | .773 [.578, .885] | *** (excellent) |
| Number of cycles per bursts (pool) | 34 | .866 [.751, .931] | *** (excellent) | Burst duration mean (pool) | 30 | .762 [.562, .879] | *** (excellent) |
| Number of burst cycles (pool) | 34 | .856 [.731, .925] | *** (excellent) | Number of cycles per bursts (all) | 30 | .761 [.559, .878] | *** (excellent) |
| Burst duration mean (pool) | 33 | .843 [.708, .919] | *** (excellent) | Burst duration SD | 30 | .752 [.539, .874] | *** (excellent) |
| Number of burst cycles (pool) | 34 | .823 [.676, .908] | *** (excellent) | Bursts per second (all) | 30 | .748 [.538, .871] | ** (good) |
| Burst duration mean (all) | 33 | .795 [.627, .893] | *** (excellent) | Number of bursts (all) | 30 | .748 [.538, .871] | ** (good) |
| Burst duration SD (pool) | 33 | .775 [.594, .882] | *** (excellent) | Number of burst cycles (pool) | 30 | .744 [.532, .869] | ** (good) |
| Number of cycles per bursts (all) | 33 | .767 [.581, .877] | *** (excellent) | Number of bursts (pool) | 30 | .712 [.481, .851] | ** (good) |
| Variance of the number of cycles per burst (pool) | 33 | .726 [.516, .854] | ** (good) | Number of cycles per bursts (pool) | 30 | .708 [.475, .849] | ** (good) |
| Burst duration SD (pool) | 33 | .717 [.502, .849] | ** (good) | Bursts per second (pool) | 30 | .708 [.475, .849] | ** (good) |
| Burst amplitude (pool) | 34 | .707 [.491, .842] | ** (good) | Burst duration SD (pool) | 30 | .691 [.450, .839] | ** (good) |
| Number of bursts (all) | 34 | .690 [.463, .832] | ** (good) | Variance of the number of cycles per burst (all) | 30 | .649 [.386, .815] | ** (good) |
| Number of bursts (pool) | 34 | .646 [.403, .805] | ** (good) | Burst band amplitude (all) | 30 | .599 [.305, .788] | * (moderate) |
| Variance of the number of cycles per burst (all) | 33 | .646 [.396, .807] | ** (good) | Variance of the number of cycles per burst (pool) | 29 | .536 [.227, .749] | * (moderate) |
| Burst band amplitude (pool) | 33 | .608 [.343, .785] | ** (good) | Burst amplitude (all) | 30 | .517 [.197, .737] | * (moderate) |
| Burst band amplitude (all) | 33 | .607 [.340, .784] | ** (good) | Burst band amplitude (pool) | 29 | .511 [.188, .736] | * (moderate) |
| Burst amplitude (all) | 33 | .597 [.326, .778] | * (moderate) | Burst amplitude (pool) | 29 | .438 [.104, .687] | * (moderate) |
| Burst PT symmetry (pool) | 33 | .592 [.320, .774] | * (moderate) | Burst PT symmetry (pool) | 30 | .334 [0.016, .614] | . (low) |
| Burst PT symmetry (all) | 33 | .382 [.068, .632] | . (low) | Burst PT symmetry (all) | 28 | .295 [0.078, .596] | . (low) |
| Burst RD symmetry (pool) | 33 | .256 [0.078, .543] | . (low) | Burst RD symmetry (pool) | 28 | .121 [0.256, .467] | ns (poor) |
| Burst RD symmetry (all) | 34 | .000 [0.334, .334] | ns (poor) | Burst RD symmetry (all) | 30 | .000 [0.355, .355] | ns (poor) |

**Note.** All = all channels; Pool = cluster-level. PT: peak-to-trough; RD: rise-decay. Significance levels are indicated as follows: $p < .05$*, $p < .01$**, $p < .001$***.

#### Table S2. Descriptive statistics and group comparisons for EEG spectral measures and alpha burst dynamics.

| Session | Variable | M (SD) YA | M (SD) OA | Statistic | ES | *p* | *p*FDR | Direction |
|---|---|---|---|---|---|---|---|---|
| T1 | iAPF | 10.55 (0.95) | 10.41 (0.93) | t(63) = 0.61 | d = 0.15 | .545 | .631 | - |
| T1 | Alpha power | 18.86 (11.68) | 16.04 (13.11) | t(65) = 0.93 | d = 0.23 | .355 | .631 | - |
| T1 | Burst duration mean (all) | 615.7 (70.7) | 635.4 (63.6) | t(61) = -1.18 | d = -0.29 | .241 | .389 | - |
| T1 | Burst duration SD (all) | 370.1 (91.0) | 379.9 (73.3) | t(58) = -0.48 | d = -0.12 | .633 | .719 | - |
| **T1** | **Bursts per second (all)** | **0.542 (0.066)** | **0.582 (0.055)** | **W = 363** | **r = -0.35** | **.014*** | **.061** | **OA > YA** |
| **T1** | **Burst band amplitude (all)** | **2.09e-06 (8.89e-07)** | **1.67e-06 (5.38e-07)** | **t(48) = 2.28** | **d = 0.58** | **.027*** | **.081** | **YA > OA** |
| T1 | Burst period (all) | 104.1 (3.3) | 103.7 (2.5) | t(55) = 0.54 | d = 0.14 | .590 | .698 | - |
| **T1** | **Burst PT symmetry (all)** | **0.503 (0.001)** | **0.504 (0.001)** | **t(51) = -2.79** | **d = -0.71** | **.007**** | **.035*** | **OA > YA** |
| T1 | Burst RD symmetry (all) | 0.499 (0.001) | 0.499 (0.001) | W = 653 | r = 0.17 | .235 | .389 | - |
| T1 | Burst amplitude (all) | 6.32e-06 (2.24e-06) | 5.60e-06 (1.58e-06) | t(53) = 1.50 | d = 0.38 | .140 | .268 | - |
| **T1** | **Number of bursts (all)** | **129.4 (15.5)** | **137.8 (14.0)** | **t(61) = -2.30** | **d = -0.57** | **.025*** | **.081** | **OA > YA** |
| **T1** | **Number of burst cycles (all)** | **777.9 (164.6)** | **865.3 (171.3)** | **t(64) = -2.12** | **d = -0.52** | **.038*** | **.098** | **OA > YA** |
| T1 | Number of cycles per burst (all) | 5.92 (0.66) | 6.14 (0.61) | t(62) = -1.38 | d = -0.34 | .174 | .317 | - |
| T1 | Variance of the number of cycles per burst (all) | 14.44 (7.21) | 14.72 (5.65) | W = 485 | r = -0.11 | .464 | .619 | - |
| **T1** | **Relative burst time (all)** | **32.67 (7.01)** | **36.47 (7.11)** | **t(64) = -2.20** | **d = -0.54** | **.032*** | **.089** | **OA > YA** |
| **T1** | **Percentage of burst samples (all)** | **33.78 (7.19)** | **37.83 (7.34)** | **t(64) = -2.27** | **d = -0.56** | **.026*** | **.081** | **OA > YA** |
| T1 | Burst duration mean (pool) | 655.4 (92.8) | 666.9 (83.7) | W = 501 | r = -0.08 | .598 | .698 | - |
| T1 | Burst duration SD (pool) | 420.0 (120.5) | 416.6 (98.0) | W = 539 | r = -0.01 | .969 | .969 | - |
| **T1** | **Bursts per second (pool)** | **0.571 (0.066)** | **0.606 (0.053)** | **t(58) = -2.44** | **d = -0.61** | **.018*** | **.068** | **OA > YA** |
| T1 | Burst band amplitude (pool) | 2.32e-06 (1.14e-06) | 1.78e-06 (7.42e-07) | W = 670 | r = 0.24 | .103 | .227 | - |
| T1 | Burst period (pool) | 103.0 (3.8) | 102.7 (2.9) | t(56) = 0.33 | d = 0.08 | .742 | .799 | - |
| T1 | Burst PT symmetry (pool) | 0.504 (0.002) | 0.504 (0.002) | W = 503 | r = -0.10 | .493 | .628 | - |
| T1 | Burst RD symmetry (pool) | 0.499 (0.002) | 0.499 (0.002) | t(62) = -0.72 | d = -0.18 | .471 | .619 | - |
| T1 | Burst amplitude (pool) | 6.72e-06 (2.89e-06) | 6.09e-06 (2.41e-06) | W = 611 | r = 0.09 | .509 | .629 | - |
| **T1** | **Number of bursts (pool)** | **136.3 (15.4)** | **143.5 (13.7)** | **t(61) = -2.00** | **d = -0.49** | **.050*** | **.123** | **OA > YA** |
| T1 | Number of burst cycles (pool) | 877.9 (197.2) | 953.2 (200.2) | t(64) = -1.55 | d = -0.38 | .127 | .253 | - |
| T1 | Number of cycles per burst (pool) | 6.37 (0.91) | 6.59 (0.94) | t(64) = -0.94 | d = -0.23 | .349 | .543 | - |
| T1 | Variance of the number of cycles per burst (pool) | 19.19 (11.15) | 18.11 (8.51) | W = 547 | r = 0.01 | .959 | .969 | - |
| T1 | Relative burst time (pool) | 36.63 (8.33) | 39.91 (8.18) | t(63) = -1.62 | d = -0.40 | .111 | .233 | - |
| T1 | Percentage of burst samples (pool) | 37.75 (8.48) | 41.28 (8.40) | t(63) = -1.71 | d = -0.42 | .093 | .216 | - |
| T2 | iAPF | 10.52 (0.84) | 10.36 (1.05) | t(61) = 0.66 | d = 0.16 | .512 | .631 | - |
| T2 | Alpha power | 18.57 (9.93) | 17.16 (13.24) | t(61) = 0.48 | d = 0.12 | .631 | .631 | - |
| **T2** | **Burst duration mean (all)** | **597.0 (45.7)** | **647.0 (68.0)** | **W = 274** | **r = -0.44** | **.003**** | **.016*** | **OA > YA** |
| **T2** | **Burst duration SD (all)** | **348.1 (64.4)** | **391.5 (78.4)** | **W = 335** | **r = -0.34** | **.019*** | **.060** | **OA > YA** |
| **T2** | **Bursts per second (all)** | **0.534 (0.057)** | **0.583 (0.045)** | **t(56) = -3.77** | **d = -0.96** | **< .001***** | **.006**** | **OA > YA** |
| T2 | Burst band amplitude (all) | 1.74e-06 (8.14e-07) | 1.76e-06 (9.22e-07) | W = 510 | r = 0.00 | 1.000 | 1.000 | - |
| T2 | Burst period (all) | 104.0 (3.5) | 104.1 (2.9) | t(56) = -0.12 | d = -0.03 | .909 | .931 | - |
| **T2** | Burst PT symmetry (all) | **0.503 (0.001)** | **0.504 (0.001)** | **t(58) = -4.27** | **d = -1.09** | **< .001***** | **.003**** | **OA > YA** |
| T2 | Burst RD symmetry (all) | 0.499 (0.001) | 0.499 (0.001) | t(54) = 1.89 | d = 0.48 | .064 | .150 | - |
| T2 | Burst amplitude (all) | 5.30e-06 (2.05e-06) | 5.82e-06 (2.50e-06) | W = 451 | r = -0.12 | .431 | .584 | - |
| **T2** | **Number of bursts (all)** | **127.5 (13.9)** | **140.4 (12.5)** | **W = 259** | **r = -0.49** | **< .001***** | **.006**** | **OA > YA** |
| **T2** | **Number of burst cycles (all)** | **751.2 (137.0)** | **880.9 (141.1)** | **t(61) = -3.73** | **d = -0.93** | **< .001***** | **.006**** | **OA > YA** |
| **T2** | Number of cycles per burst (all) | **5.81 (0.54)** | **6.22 (0.61)** | **t(62) = -2.88** | **d = -0.72** | **.005**** | **.025*** | **OA > YA** |
| **T2** | Variance of the number of cycles per burst (all) | **12.48 (5.06)** | **15.43 (6.16)** | **W = 351** | **r = -0.31** | **.033*** | **.087** | **OA > YA** |
| **T2** | **Relative burst time (all)** | **31.53 (5.60)** | **36.67 (6.11)** | **t(62) = -3.51** | **d = -0.88** | **< .001***** | **.006**** | **OA > YA** |

| | | | | | | | | |
|---|---|---|---|---|---|---|---|---|
| **T2** | **Percentage of burst samples (all)** | **32.58 (5.72)** | **38.07 (6.35)** | **t(62) = -3.64** | **d = -0.90** | **< .001*** ** | **.006** ** | **OA > YA** |
| T2 | Burst mean duration (pool) | 639.8 (72.2) | 682.7 (98.0) | W = 378 | r = -0.26 | .077 | .161 | - |
| T2 | Burst duration SD (pool) | 393.5 (89.6) | 433.2 (113.4) | W = 400 | r = -0.22 | .141 | .246 | - |
| **T2** | **Bursts per second (pool)** | **0.575 (0.055)** | **0.606 (0.047)** | **t(57) = -2.39** | **d = -0.61** | **.020*** | **.060** | **OA > YA** |
| T2 | Burst band amplitude (pool) | 1.79e-06 (8.98e-07) | 1.78e-06 (1.01e-06) | W = 489 | r = 0.02 | .888 | .931 | - |
| T2 | Burst period (pool) | 102.7 (3.7) | 103.2 (3.5) | t(60) = -0.58 | d = -0.15 | .562 | .715 | - |
| T2 | Burst PT symmetry (pool) | 0.504 (0.002) | 0.504 (0.002) | t(61) = -1.69 | d = -0.42 | .095 | .174 | - |
| T2 | Burst RD symmetry (pool) | 0.499 (0.001) | 0.499 (0.001) | t(59) = -0.39 | d = -0.10 | .697 | .808 | - |
| T2 | Burst amplitude (pool) | 5.27e-06 (2.24e-06) | 6.15e-06 (3.13e-06) | W = 415 | r = -0.16 | .285 | .399 | - |
| **T2** | **Number of bursts (pool)** | **137.2 (13.5)** | **145.9 (13.3)** | **t(61) = -2.58** | **d = -0.65** | **.012*** | **.052** | **OA > YA** |
| **T2** | **Number of burst cycles (pool)** | **864.4 (165.2)** | **970.3 (184.8)** | **t(62) = -2.42** | **d = -0.60** | **.018*** | **.060** | **OA > YA** |
| T2 | Number of cycles per burst (pool) | 6.24 (0.73) | 6.61 (0.89) | t(62) = -1.83 | d = -0.45 | .072 | .160 | - |
| T2 | Variance of the number of cycles per burst (pool) | 15.41 (6.08) | 18.31 (8.22) | W = 389 | r = -0.19 | .209 | .314 | - |
| **T2** | **Relative burst time (pool)** | **36.02 (6.75)** | **40.18 (7.91)** | **t(62) = -2.27** | **d = -0.56** | **.027*** | **.074** | **OA > YA** |
| **T2** | **Percentage of burst samples (pool)** | **37.08 (6.90)** | **41.60 (8.17)** | **t(62) = -2.40** | **d = -0.60** | **.019*** | **.060** | **OA > YA** |

**Note.** Values are presented as *M* (SD). Extreme values (> 3 SD from the group mean, computed separately within each group and variable) were replaced with NA prior to statistical analyses. Group differences were assessed using independent-samples *t*-tests (two-tailed) or Wilcoxon rank-sum tests depending on normality. Effect sizes (ES) are reported as Cohen's *d* (Welch) or rank-biserial *r* (Wilcoxon). *p*FDR = *p*-value corrected using Benjamini-Hochberg false discovery rate procedure independently within each group and session. YA = younger adults; OA = older adults. All = all channels; Pool = cluster-level. PT: peak-to-trough; RD: rise-decay. Significance levels are indicated as follows: $p < .05$*, $p < .01$**, $p < .001$***.

**Table S3. Partial Spearman correlations controlling for years of education and the respective subjective covariate (arousal, boredom, or richness of experience).**

When controlling for arousal revealed nominal positive associations between rTE2 and alpha burst rate in older adults ($\rho = .41$, $p = .023$, $p$FDR = .300 and $\rho = .39$, $p = .029$, $p$FDR = .300) and with cycles per burst variability ($\rho = .37$, $p = .046$, $p$FDR = .300). Controlling for boredom revealed a nominal positive association between rTE2 and burst duration in older adults ($\rho = .37$, $p = .045$, $p$FDR = .432). Controlling for richness revealed a nominal negative association between rTE1 and the number of alpha bursts in younger adults ($\rho = -.39$, $p = .047$, $p$FDR = .875). However, none of these associations survived FDR correction.

| Covariate | Group | Session | Temporal measure | EEG metric | n | *ρ* | *p* | *p*FDR |
|---|---|---|---|---|---|---|---|---|
| Arousal | OA | T2 | rTE2 | Bursts per second (pool) | 34 | .41 | .023* | .300 |
| Arousal | OA | T2 | rTE2 | Bursts per second (all) | 34 | .39 | .029* | .300 |
| Arousal | OA | T2 | rTE2 | Variance of the number of cycles per burst (pool) | 33 | .36 | .046* | .300 |
| Boredom | OA | T2 | rTE2 | Burst mean duration (all) | 33 | .36 | .045* | .432 |
| Richness | YA | T1 | rTE1 | Number of bursts (pool) | 29 | -.38 | .047* | .875 |

**Note.** Only associations reaching nominal significance ($p < .05$) are reported. None of the associations survived FDR correction. YA = younger adults; OA = older adults; rTE1/rTE2 = retrospective time estimate at Session 1 and 2. All = all channels; Pool = cluster-level. Significance levels are indicated as follows: $p < .05$*, $p < .01$**, $p < .001$***.

**Table S4. Associations between temporal measures and neuropsychological measures across age groups.**

Nominal correlations between neuropsychological measures and temporal measures (rTE1, rTE2, duration ratios across video clips) in older (OA) and younger adults (YA). Only associations reaching $p$ < .05 are reported. Correlations were computed using Pearson or Spearman methods depending on normality assumptions. *p*FDR values were adjusted using the Benjamini-Hochberg procedure within each group. No significant associations were observed after FDR correction in either group. In younger adults, rTE2 was positively associated with autobiographical memory richness (ART: $\rho$ = .52, $p$ = .005) and working memory (N-Back: $\rho$ = .46, $p$ = .013). rTE2 was also associated with internal attentional orientation (ASQ Internal: $\rho$ = .40, $p$ = .037) and chronotype ($\rho$ = .38, $p$ = .045). rTE1 was positively associated with perceived recent changes ($\rho$ = .52, $p$ = .004), negative past temporal perspective (ZTPI Negative Past: $r$ = .40, $p$ = .030), and subjective temporality ($r$ = .41, $p$ = .026). In older adults, rTE1 was positively associated with negative past temporal perspective (ZTPI Negative Past: $r$ = .35, $p$ = .039), rTE2 was positively associated with subjective temporality ($\rho$ = .41, $p$ = .021) and negatively associated with fatalistic present temporal perspective (ZTPI Fatalistic Present: $\rho$ = -.36, $p$ = .044). For the video-based duration estimation task, in older adults, cognitive impulsivity (BIS-11) was positively associated with duration ratios across all clip durations (2-s to 16-s; *r*s and *ρ*s = .36-.55, *p*s < .038, *p*FDR < .048), as was trait impulsivity (BIS-11 total score; *r*s and *ρ*s = .40-.58, *p*s < .022, *p*FDR < .028) and planning difficulties (BIS-11 planning; $\rho$ = .48, $p$ = .005, *p*FDR = .050). Working memory was also negatively associated with the 8-s low-eventfulness ratio (Forward digit span: $\rho$ = -.50, $p$ = .004, *p*FDR = .040).

| Group | Temporal measure | Neuropsychological measure | n | r/ρ | p | pFDR |
|---|---|---|---|---|---|---|
| OA | rTE1 | ZTPI Negative Past | 35 | .35 | .039* | .968 |
| OA | rTE2 | Temporality | 34 | .41 | .021* | .580 |
| OA | rTE2 | ZTPI Present Fatalist | 34 | -.36 | .044* | .580 |
| OA | ASQ External Orientation | ratio_2000_low | 34 | .44 | .011* | .052 |
| OA | ASQ External Orientation | ratio_4000_low | 34 | .43 | .012* | .052 |
| OA | ASQ External Orientation | ratio_4000_high | 34 | .42 | .016* | .052 |
| OA | ASQ External Orientation | ratio_8000_high | 34 | .36 | .039* | .079 |
| OA | ASQ External Orientation | ratio_16000_low | 35 | .34 | .048* | .079 |
| OA | ASQ External Orientation | ratio_2000_high | 33 | .35 | .048* | .079 |
| **OA** | **BIS-11 Planning difficulties** | **ratio_8000_high** | **34** | **.48** | **.005**** | **.050*** |
| OA | BIS-11 Planning difficulties | ratio_16000_high | 34 | .40 | .022* | .098 |
| OA | BIS-11 Planning difficulties | ratio_16000_low | 35 | .37 | .031* | .098 |
| OA | BIS-11 Planning difficulties | ratio_8000_low | 34 | .36 | .039* | .098 |
| **OA** | **BIS-11 Cognitive Impulsivity** | **ratio_8000_high** | **34** | **.55** | **<.001***** | **.009**** |
| **OA** | **BIS-11 Cognitive Impulsivity** | **ratio_8000_low** | **34** | **.48** | **.005**** | **.022*** |
| **OA** | **BIS-11 Cognitive Impulsivity** | **ratio_2000_low** | **34** | **.45** | **.008**** | **.022*** |
| **OA** | **BIS-11 Cognitive Impulsivity** | **ratio_4000_high** | **34** | **.44** | **.011*** | **.022*** |
| **OA** | **BIS-11 Cognitive Impulsivity** | **ratio_4000_low** | **34** | **.44** | **.011*** | **.022*** |
| **OA** | **BIS-11 Cognitive Impulsivity** | **ratio_16000_low** | **35** | **.36** | **.037*** | **.047*** |

| | | | | | | |
|---|---|---|---|---|---|---|
| OA | **BIS-11 Cognitive Impulsivity** | **ratio_2000_high** | **33** | **.37** | **.037*** | **.047*** |
| OA | **BIS-11 Cognitive Impulsivity** | **ratio_16000_high** | **34** | **.36** | **.038*** | **.047*** |
| OA | BIS-11 Motor Impulsivity | ratio_8000_high | 33 | .46 | .008** | .078 |
| OA | BIS-11 Motor Impulsivity | ratio_4000_high | 33 | .40 | .024* | .078 |
| OA | BIS-11 Motor Impulsivity | ratio_4000_low | 33 | .39 | .029* | .078 |
| OA | BIS-11 Motor Impulsivity | ratio_2000_high | 32 | .39 | .031* | .078 |
| OA | BIS-11 Motor Impulsivity | ratio_2000_low | 33 | .35 | .050* | .099 |
| OA | **BIS-11 Total Score** | **ratio_8000_high** | **34** | **.58** | **<.001***** | **.004**** |
| OA | **BIS-11 Total Score** | **ratio_8000_low** | **34** | **.50** | **.003**** | **.017*** |
| OA | **BIS-11 Total Score** | **ratio_16000_high** | **34** | **.47** | **.006**** | **.020*** |
| OA | **BIS-11 Total Score** | **ratio_4000_high** | **34** | **.44** | **.010**** | **.024*** |
| OA | **BIS-11 Total Score** | **ratio_4000_low** | **34** | **.42** | **.014*** | **.027*** |
| OA | **BIS-11 Total Score** | **ratio_2000_low** | **34** | **.41** | **.019*** | **.027*** |
| OA | **BIS-11 Total Score** | **ratio_16000_low** | **35** | **.40** | **.020*** | **.027*** |
| OA | **BIS-11 Total Score** | **ratio_2000_high** | **33** | **.40** | **.022*** | **.027*** |
| OA | CVLT delayed cued recall | ratio_32000_high | 35 | .38 | .027* | .266 |
| OA | CVLT Immediate cued recall | ratio_32000_high | 35 | .41 | .016* | .162 |
| OA | **Forward digit span** | **ratio_8000_low** | **33** | **-.50** | **.004**** | **.040*** |
| OA | Forward digit span | ratio_2000_low | 33 | -.43 | .015* | .053 |
| OA | Forward digit span | ratio_4000_high | 33 | -.42 | .017* | .053 |
| OA | Forward digit span | ratio_16000_low | 34 | -.39 | .024* | .053 |
| OA | Forward digit span | ratio_8000_high | 33 | -.39 | .029* | .053 |
| OA | Forward digit span | ratio_2000_high | 32 | -.39 | .032* | .053 |
| OA | Chronotype (MSFsc) | ratio_16000_high | 34 | .41 | .019* | .188 |
| OA | ZTPI Future | ratio_8000_high | 34 | -.40 | .022* | .225 |
| YA | rTE1 | Recent changes | 30 | .52 | .004** | .131 |
| YA | rTE1 | Temporality | 30 | .41 | .026* | .371 |
| YA | rTE1 | ZTPI Negative Past | 30 | .40 | .030* | .371 |
| YA | rTE2 | Autobiographical Recollection Test | 29 | .52 | .005** | .203 |
| YA | rTE2 | N-Back | 30 | .46 | .013* | .243 |
| YA | rTE2 | ASQ Internal Orientation | 30 | .40 | .037* | .330 |
| YA | rTE2 | Chronotype (MSFsc) | 30 | .38 | .045* | .330 |
| YA | GDS | ratio_32000_high | 30 | .50 | .006** | .058 |
| YA | GDS | ratio_32000_low | 30 | .43 | .020* | .095 |
| YA | GDS | ratio_16000_low | 29 | .41 | .028* | .095 |
| YA | HADS Anxiety | ratio_8000_high | 29 | .39 | .039* | .199 |

**Note.** *r* and *ρ* = correlation coefficient; *p* = nominal *p*-value; *p*FDR = FDR-corrected *p*-value. YA = Younger Adults; OA = Older Adults. Significance levels are indicated as follows: $p < .05$*, $p < .01$**, $p < .001$***.
